# Role of Mutual Information Profile Shifts in Assessing the Pathogenicity of Mutations on Protein Functions: The Case of Pyrin Variants Associated with Familial Mediterranean Fever

**DOI:** 10.1101/2024.04.07.588500

**Authors:** Aysima Hacisuleyman, Ahmet Gul, Burak Erman

**Affiliations:** Department of Computational Biology, University of Lausanne, CH-1015 Lausanne, Switzerland; Division of Rheumatology, Department of Internal Medicine, Istanbul University, Istanbul Faculty of Medicine, Istanbul, Turkey; Chemical and Biological Engineering, Koc University, Istanbul, Turkey

## Abstract

This paper presents a novel method to assess the pathogenicity of Pyrin protein mutations by using mutual information (MI) as a measure to quantify the correlation between residue motions or fluctuations and associated changes affecting the phenotype. The concept of MI profile shift is presented to quantify changes in MI upon mutation, revealing insights into residue-residue interactions at critical positions. We apply this method to the Pyrin protein variants, which are associated with an autosomal recessively inherited disease called familial Mediterranean fever (FMF) since the available tools do not help predict the pathogenicity of the most penetrant variants. We demonstrate the utility of MI profile shifts in assessing the effects of mutations on protein stability, function, and disease phenotype. The importance of MI shifts, for the pyrin example the negative shifts, as indicators of severe functional effects is emphasized, along with exploring potential compensatory mechanisms indicated by positive MI shifts, which are otherwise random and inconsequential. The paper also discusses challenges in relating MI profile changes to disease severity and advocates for comprehensive analysis considering genetic, environmental, and stochastic factors. Overall, this study provides insights into the molecular mechanisms underlying the pathogenesis of FMF and offers a framework for identifying potential therapeutic targets based on MI profile changes induced by mutations.

## Introduction

The specific purpose of this paper is to develop a method for assessing the effects of missense variations in amino acid sequence (henceforth mutation) on protein dynamics, and interactions and predicting their pathogenicity, with a particular focus on understanding how changes in mutual information (MI) contribute to the pathogenicity of a mutation associated with a particular phenotype. Mutual Information (MI) quantifies the degree of information shared between two random variables. These variables can represent fluctuations in a variety of properties, including biological, chemical, electromagnetic, or mechanical attributes of residues. In the context of protein dynamics, we focus on fluctuations in the positions of residues over time. Mutual information (MI) between two residues serves as a measure of the statistical dependence or correlation between their respective motions or fluctuations. More specifically, MI quantifies the amount of information shared between the behaviors of the two residues, reflecting how knowing the state or behavior of one residue informs us about the state or behavior of the other residue, and vice versa. In essence, MI provides insights into how changes in one residue may influence the behavior of the other^1^. When the motions of two residues are highly correlated or coupled, they will exhibit a high MI, indicating that changes in one residue are likely to be associated with changes in the other residue. Conversely, low MI suggests a weaker correlation, where changes in one residue may have less predictable effects on the behavior of the other. Therefore, MI offers valuable insights into the dynamic interplay between residues within a protein and aids in understanding how structural and functional changes propagate through the protein structure^2, 3^.

MI can be calculated based on the probability distributions of the fluctuations or motions of the residues. It provides a quantitative measure of the strength of the relationship between residues in terms of their dynamics, allowing one to identify residues that tend to move together or influence each other’s motions. Mathematically, MI is defined as

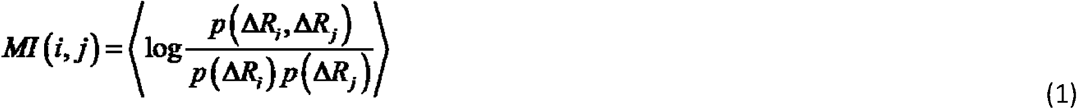

where, *p* (Δ*R*_*i*_, Δ*R*_*j*_) is the joint probability of the fluctuations of two residues i and j. *p* (Δ*R*_*i*_) and *p* (Δ*R*_*j*_) are the singlet probabilities. The variables will be functions of time, and the angular brackets denote averaging over time.

In the context of our research on Pyrin and its mutants, calculating MI between pairs of residues allows us to assess how these mutations affect the correlated motions of residues within the protein structure. Comparing MI values between the wild-type and mutant proteins enables us to quantify the impact of mutations on residue-residue interactions and infer how these changes may influence protein dynamics, function, and disease. For this purpose, we introduce the MI profile shift, Δ*MI* (*i*) which we define as

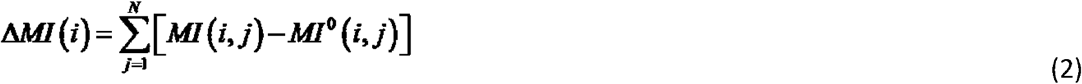

where, *MI* (*i, j*) defined in Eq. (1) represents the mutual information between the fluctuations of residue i and residue j in the mutated protein. MI is a non-negative quantity that captures the statistical dependence between two variables. In our case, it quantifies the information shared between the fluctuations of two amino acid residues, indicating how much knowing the fluctuation of one residue tells us about the fluctuation of the other. *MI* ^0^ (*i, j*) is the mutual information between residues i and j in the wild type, WT, structure, and the summation is for all residues of the protein. Δ*MI* (*i*) represents a shift in the profile of mutual information values across the protein, with residue i serving as a focal point for the analysis. This measure is meaningful because it quantifies the change in mutual information between residue i and the rest of the protein upon mutation. Alternatively, Δ*MI* (*i*) may be viewed as the coupling of residue i, i.e., the statistical coupling, to the rest of the protein. Positive Δ*MI* (*i*) signifies an increase in the net coupling between residue i and the rest of the protein. The mutation might have introduced new interactions or strengthened existing ones, leading to a more interconnected residue i. Negative Δ*MI* (*i*) indicates a decrease in the net coupling. The mutation might have disrupted communication pathways between residue i and other residues, weakening its overall connection to the protein network. Zero Δ*MI* (*i*)suggests that the mutation has resulted in no net change in the coupling of residue i. The positive and negative effects on individual interactions with other residues might have canceled each other out, leaving the overall communication of residue i with the protein relatively unchanged. By comparing this change between the wild type and the mutant, we can infer how the mutation affects the residue’s correlation with other residues. However, the application of MI profile shift can go further than simple correlation analysis^4, 5^. Proteins can be represented as networks or graphs, where residues are nodes and interactions between residues are edges. Changes in mutual information could potentially correlate with changes in graph properties such as connectivity, centrality, or modularity^6, 7^. For instance, a mutation that disrupts interactions between residues might lead to changes in network connectivity or community structure. Investigating these relationships could provide insights into how mutations affect the overall structural and functional properties of proteins^8^. In a protein interaction network, connectivity refers to the degree to which residues are connected. Mutual information captures the degree of correlation or interaction between residues. A mutation that disrupts interactions between residues could lead to a decrease in connectivity, as fewer interactions are maintained in the mutated protein compared to the wild type. Centrality measures the importance or influence of nodes in a network. Residues with high centrality are often critical for maintaining the overall structure or function of the protein. Changes in mutual information could affect the centrality of residues by altering their role in mediating interactions within the protein. A mutation that disrupts interactions involving central residues could lead to a decrease in their centrality and potentially affect the stability or function of the protein^9-12^. Modularity refers to the presence of densely connected groups of nodes, or modules, within a network. In proteins, modules can represent functional units or structural domains. Changes in mutual information could affect modularity by altering the interactions between residues within and between modules or domains^5, 13^. A mutation that disrupts interactions within a module could lead to its fragmentation or reorganization, affecting the overall modular infrastructure of the protein.

However, mutations may strengthen some interactions in the mutant protein that result from a compensatory mechanism ^14-16^. For example, if a mutation disrupts one set of interactions crucial for protein stability or function, the protein might adapt by strengthening alternative interactions to maintain its structure or activity. Understanding which interactions are strengthened and how they contribute to the overall function is important for understanding the functional consequences of mutations.

Mutations can also induce allosteric effects, where changes in one part of the protein propagate to distant regions, affecting their dynamics and interactions^17-19^. Understanding these allosteric networks is essential for understanding the mechanisms by which mutations impact protein function and regulation.

By investigating how changes in mutual information correlate with changes in these graph properties, we can gain insights into how mutations affect the structural and functional properties of proteins at a systems level. This approach allows for a more comprehensive understanding of the impact of mutations on protein dynamics and interactions, which is crucial for elucidating disease mechanisms and guiding therapeutic interventions.

However, relating the severity of a disease to the amount of MI profile shift due to mutations may not be straightforward. Firstly, the relationship between the impact of a mutation and the severity of a disease phenotype can be influenced by various factors beyond changes in MI alone. Exploring this relationship could provide valuable insights into the molecular mechanisms underlying disease pathogenesis. The specific effects of mutations on protein structure, function, and interactions can vary widely, depending on the amino acid changes and their locations within the protein. Even mutations affecting different domains of the same protein can lead to different structural, functional, and phenotypic consequences. Therefore, it is essential to consider the specific details of each mutation when comparing their impacts on MI and disease phenotype and its severity. Molecular dynamics analysis appears to be the most suitable method to pursue this. Secondly, the effects of mutations on protein dynamics and interactions can be context-dependent, influenced by factors such as the cellular environment, protein conformational changes, post-translational modifications, and protein-protein interactions. These factors can modulate the relationship between changes in MI and the pathogenicity of mutations resulting in certain disease phenotypes. Genetic modifiers and the individual genetic background of patients can influence the disease severity. Other genetic variations within the affected individual or population can affect the interactions of other proteins with the mutated protein, modulating its phenotypic expression. Therefore, the relationship between changes in MI and disease severity may be influenced by genetic factors beyond the mutated protein itself. In addition, disease severity is often determined by complex interactions between genetic, environmental, and stochastic factors. While changes in MI induced by mutations can contribute to disease pathogenesis, they are unlikely to be the sole determinant of disease severity. Other factors, such as the cumulative effects of multiple mutations, environmental triggers, and patient-specific factors, also play critical roles in shaping the disease phenotype.

We herein investigated the potential of MI profile changes using the example of the Pyrin mutations associated with a monogenic autoinflammatory disease, Familial Mediterranean Fever (FMF). Pyrin is one of the critical inflammasome components, involved in the recognition of various pathogens and damage-associated molecular patterns, which results in the activation of caspase 1 for the proteolytic maturation of IL-1beta and IL-18. Pyrin, like many proteins, has likely evolved to adopt a specific structural configuration that optimizes its function within the cellular context. This optimized structure allows Pyrin to perform its desired functions efficiently and accurately. Any deviation from this native structural landscape, such as the introduction of new intramolecular interactions due to mutations or loss of existing ones, is likely to be detrimental to protein function. These deviations can disrupt the finely tuned balance of interactions within the protein, leading to functional deficits or loss of activity. While it is theoretically possible for certain mutations to introduce new interactions that improve protein properties, such occurrences are rare and highly unlikely given the precision of evolutionary selection pressures acting on protein structure and function. Therefore, deviations from the native structural features of Pyrin are generally expected to have negative consequences, highlighting the importance of preserving the integrity of protein structure for maintaining proper cellular function^20^.

FMF is an autosomal recessively inherited disease characterized by self-limited inflammatory attacks lasting 12-72 hours, mainly involving serosal membranes and joints. FMF starts during childhood and is common in particular ethnic groups, including Jews, Armenians, Arabs, and Turks living in the Eastern Mediterranean countries. The patients experience recurrent episodes of peritonitis, pleuritis, arthritis, or erysipelas-like erythema, and up to 20-30% of them may develop AA amyloidosis due to uncontrolled inflammatory response. This typical phenotype of FMF has been associated with mutations in the C-terminal SPRY domain of the Pyrin, which is encoded by exon 10 of the *MEFV* gene. On the other hand, mutations occurring in other modular domains of Pyrin, encoded by exons 2, 3, 5, or 8 may result in various autoinflammatory phenotypes other than FMF. Among the SPRY domain mutations associated with FMF phenotype, M694V is the most penetrant one and is associated with a more severe disease course. The M680I is the second most penetrant exon 10 variant, and both mutations of the SPRY domain are associated with a higher inflammatory response and the most serious complication of AA amyloidosis ^21-23^. On the other hand, starting from the identification of the M694V and M680I mutations as the main pathogenic variations leading to the FMF phenotype, all of the computational tools used to predict the pathogenicity of these missense mutations were unhelpful. Software such as SIFT (Sorting Intolerant From Tolerant) ^24^, PolyPhen-2 (Polymorphism Phenotyping v2) ^25^, PROVEAN (Protein Variation Effect Analyzer) ^26^,SAAMBE-3D^27^, Mutabind2^28^, Dynamut2^29^, mCSM-PPI2^30^, and Varity^31^, are frequently used to predict the pathogenicity of an amino acid substitution by affecting its function using the data of sequence homology, the physical properties of amino acids, structural and comparative evolutionary considerations. However, these predictions are not always successful, and the substitution of Methionine to Isoleucine at position 680 or Methionine to a Valine at position 694 in these most penetrant mutations associated with FMF phenotype were suggested as tolerant, benign, or neutral, except for the predictions by PolyPhen-2, SAAMBE-3D, Mutabind2, DynaMut2, mCSMPPI2 and Varity for M694V, which was predicted as probably damaging (Table 1). However, the predictions for the other missense variations did not match with their clinical significance, such as highly pathogenic M680I was predicted by PolyPhen-2 as benign, and the likely pathogenic mutation K695R associated with a milder phenotype was predicted as possibly damaging with scores similar to M694V (Table 1).

**Table 1.**
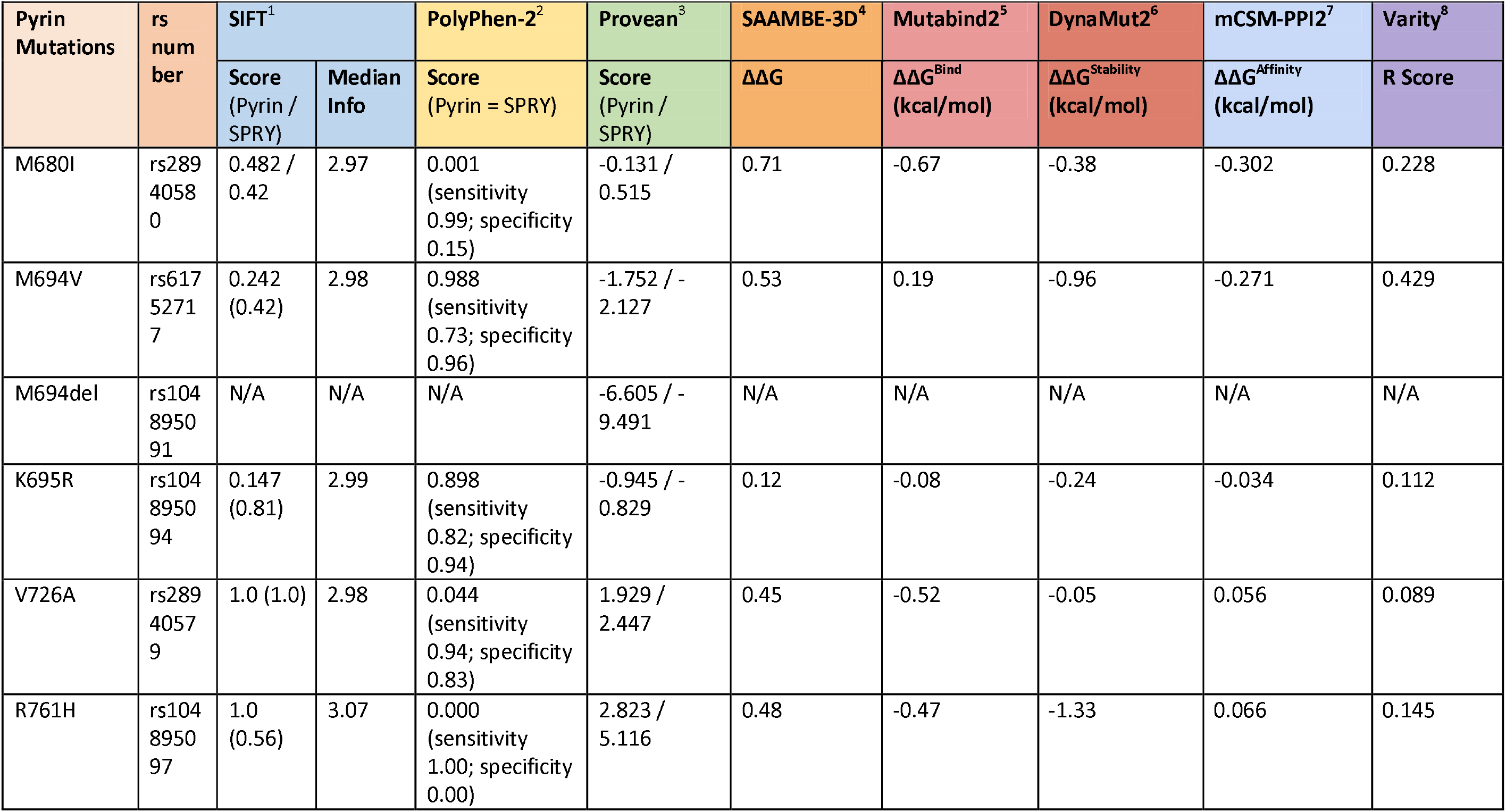
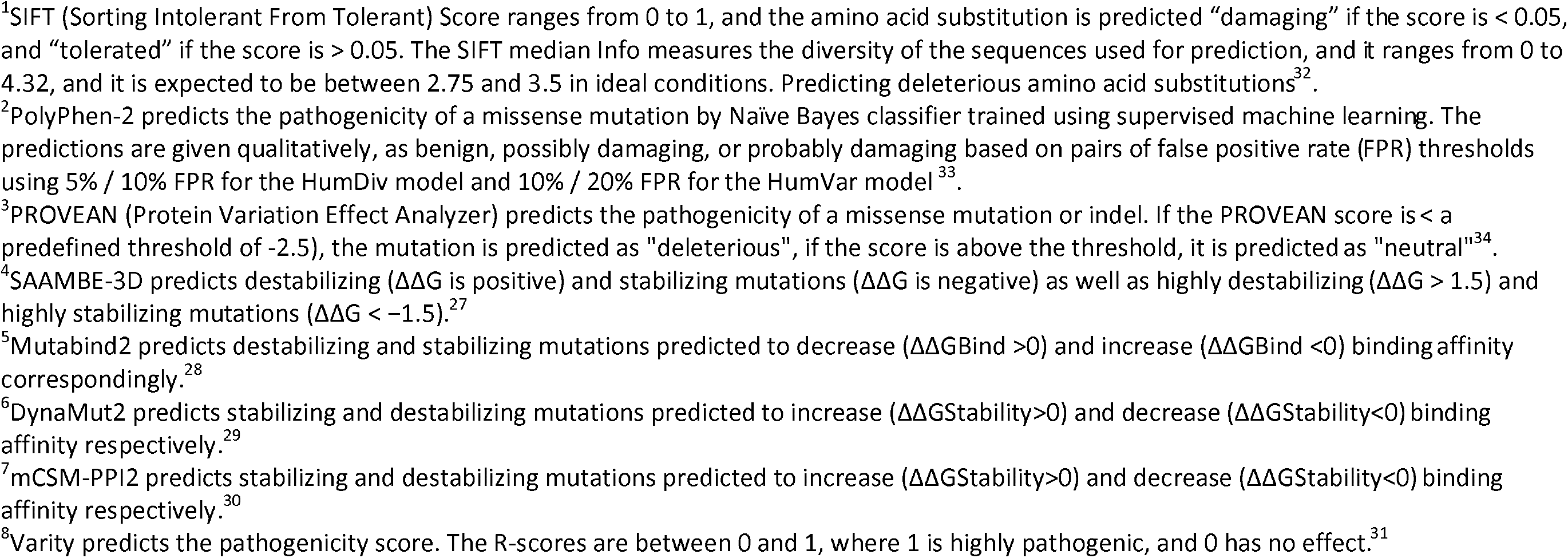
Predictions for the pathogenicity of Pyrin mutations by the SIFT, Polyphen-2, Provean, SAAMBE-3D, Mutabind2, Dynamut2, mCSM-PPI2, and the Varity servers. The scores and predictions were calculated for both the whole Pyrin protein and the SPRY domain containing the residues between 586 and 781 as used in the MD analyses.

The Alpha Missense^35^ program, which is adapted from AlphaFold2^36^, provides a database of predictions for all possible single amino acid substitutions in the human proteome as a resource to the community. However, it includes classified variants as likely pathogenic and likely benign. The retrieved data for the MEFV gene do not include the pathogenic variants we investigated in the current study. Therefore, Table 1 does not include any information about the Alpha Missense^35, 37^.

Our study employed extensive molecular dynamics analysis to investigate the shifts in mutual information profiles between the wild type and five SPRY domain mutations and one deletion (3 pathogenic mutations M694V, M680I, V726A, and 2 likely pathogenic mutations K695R, R761H; and one deletion variant M694del) associated with FMF phenotype but with varying degrees of disease severity. We also performed steered molecular dynamics (SMD)^38^ simulations to investigate the change in the binding affinity of the wild-type and six variants of the SPRY domain to actin. The exceptional deletion variant of M694del was used as an example of extreme disruption of the structure and function relationship, which results in an autosomal dominantly inherited severe FMF phenotype ^39^.

The SPRY domain has a shallow cavity for binding putative protein ligands. Although several proteins, including caspase 1, beta-2 microglobulin, and beta-actin, have been shown to be interacting with the SPRY domain, none of them was documented as the ligand being affected by the pathogenic variants and responsible for the pathogenesis of FMF^40^. We, therefore, selected beta-actin for the study since the cytoplasmic expression of Pyrin colocalizes with the expression of actin, and changes in the actin cytoskeleton through inactivation of RhoA proteins result in activation of Pyrin inflammasome ^41, 42^.

In the following sections, we provide detailed computational methods and present results through graphical representations for each mutation, highlighting discernible patterns that elucidate the relationship between the magnitude of mutual information profile shifts and mutation-associated disease severity. Finally, we offer a comprehensive discussion on the rationale behind our approach, along with insights obtained from clinical observations, to provide a comprehensive understanding of the implications of our findings.

## Methods

### System Preparation

#### MD simulations

The crystal structure of the Pyrin protein C-terminal PRY/SPRY domain, with PDB id 2WL1, was used for the simulations ^43^. The crystal waters were retained. The system was solvated in a TIP3P water box ^44^ with a buffer size of 15 Å. To neutralize the system, NaCl ions were introduced, resulting in a salt concentration of 0.15 mol/L. Charmm36 force field^45, 46^ was used to parametrize the system. A distance cutoff of 12 Å was applied to short-range, non-bonded interactions, and 10.0 Å for the smothering functions. Long-range electrostatics were managed with the particle-mesh Ewald (PME) method ^47^. The systems were prepared by using the QwikMD plugin^48^ of VMD ^49^. Prior to MD simulations, all systems underwent a 2000-step energy minimization. The time step of 2 fs was used for integration. Annealing was performed after the minimization to increase the temperature of the system from 60 K to 300 K while keeping the pressure at 1 atm using Nosé-Hoover Langevin piston ^50^. Equilibration was performed following the annealing step for 1 ns without applying any constraints on the system. The temperature was maintained at 300 K using Langevin dynamics. Finally, MD simulations were performed under NpT ensemble for 100 ns and 3 replicas were run for each system, making the total simulation time 300 ns/system. All results reported are based on the total 300 ns/system trajectory lengths.

#### Construction of probability distribution functions for pair distances, pdf

All calculations in the present study are based on alpha carbons of residues. Calculations for other atoms can be made from the all-atom trajectories from the MD simulations. The instantaneous position of each alpha carbon is recorded from which instantaneous distances between pairs are calculated. Distances for the pair of interest are arranged in increasing order and divided into 20 bins and the probability distribution function for the full range is evaluated.

#### SMD simulations

To generate the Pyrin domain in complex with Actin, we utilized Alphafold2^36^. The best model was selected from the results and the system was prepared in the same way as explained in the previous section. SMD ^38^ simulations were performed after minimization, annealing, and equilibration runs. It’s worth noting that the preparation steps, the minimization, annealing, and equilibration protocols prior to running the SMD simulations mirrored those described for the MD simulations.

The SMD was performed with explicit solvent using the TIP3 water model in the NpT ensemble. The temperature was maintained at 300 K using Langevin dynamics and the pressure was kept at 1 atm using Nosé-Hoover Langevin piston. The short- and long-range interaction cutoffs and the methods to manage them were kept the same as the MD simulations. The time step of integration was chosen to be 2 fs for all simulations. Constant velocity pulling was applied with a pulling speed of 2.5 Å/ns and a harmonic constraint force of 7.0 kcal/mol/Å^2^. In this step, SMD was employed by harmonically restraining the position of the A331 residue of the Pyrin and moving the restraint residues G719 and P754 of Actin, with constant velocity in the axis defined by the two groups of restrained residues. 10 replicates of SMD simulations were performed for each system. Figure 1 depicts how the SMD simulations are conducted.

**Figure 1.**
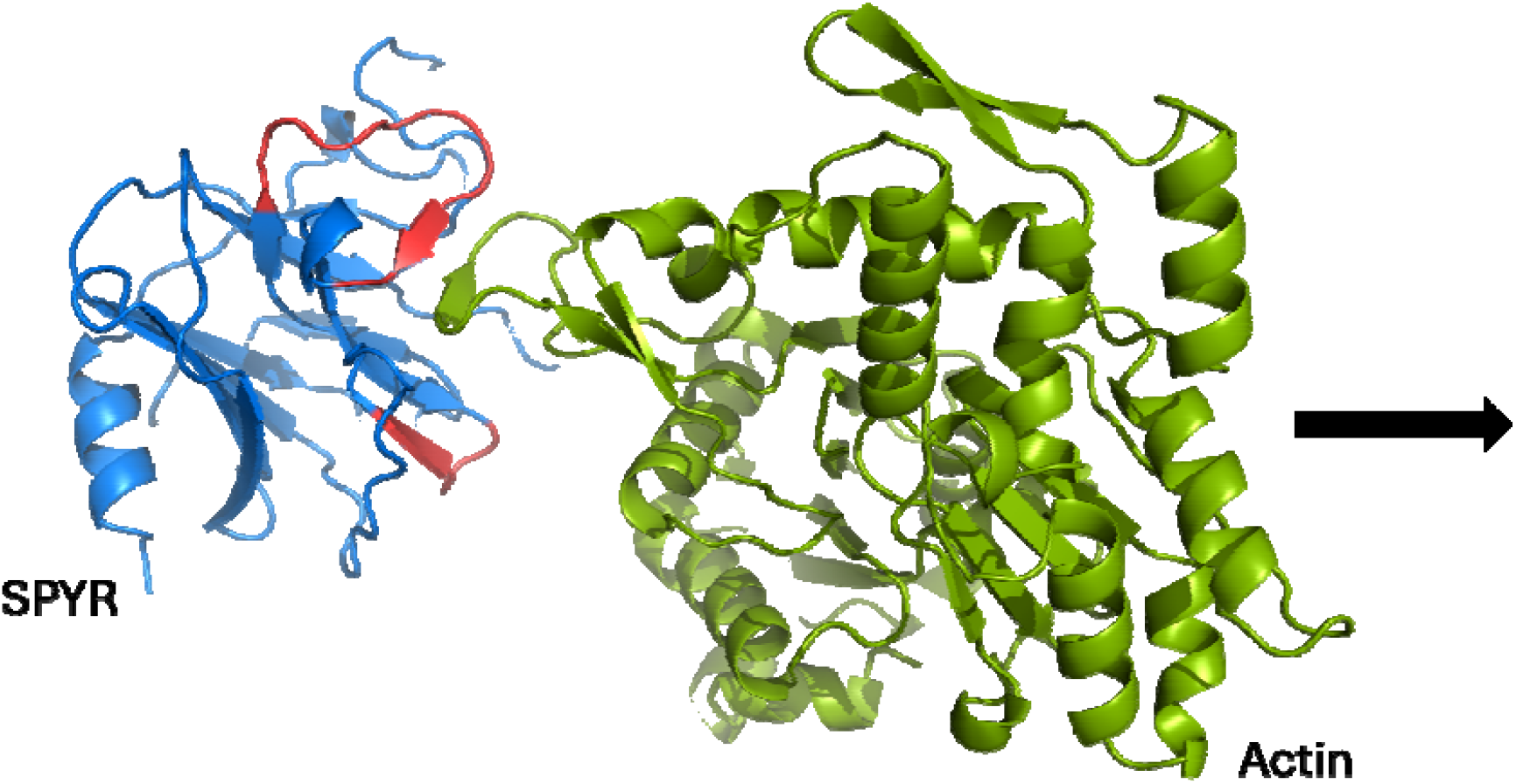
SPYR domain in blue and Actin in green shown in cartoon representation. The loops on SPYR where actin binds are colored in red. The pulling direction for the SMD simulations were shown in the figure.

#### Solvent Accessible Surface Area (SASA) measurement of the binding cavity

To quantify the structural differences of the cavity where Actin binds on the Pyrin-SPRY domain between the WT and its mutants, we measured the solvent-accessible surface area(SASA) of the cavity between loops (residue 671-683 and 695-698), where actin binds, by using the measure sasa function of VMD.

#### SMD Force, Distance and K_D_ calculations

To calculate and compare the binding affinity of Actin with the WT and mutants of Pyrin, we performed SMD simulations. The binding affinity is correlated with the rupture force. The force vectors were obtained from the simulation log file, and they were plotted against the pulling distance. The cumulative work was obtained by calculating the area under the force vs distance curve. The rupture distances were detected by determining the time when the second derivative of the cumulative work became zero, and the K_D_ values were obtained from the following formula^51^

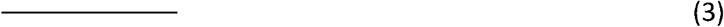

where the cumulative work is in kJ/mol, k is the Boltzmann constant, and T is the temperature. We considered kT as 2.479 kJ/mol at 300 K ^51^.

#### Mutual information profiles

The definition of mutual information is based on the instantaneous fluctuation Δ*R*_*i*_ (*t*) of residue i at time t from molecular dynamics simulations. In Cartesian coordinates,

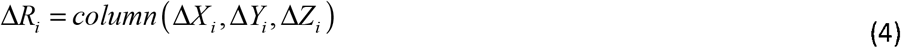

and the joint fluctuation of i and j is

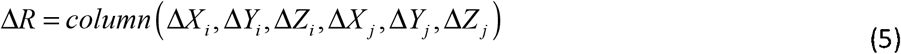

Where **Δ*R*** constitutes the state vector defined for each time point of the MD simulations.

We considered only the alpha carbons in the calculations and used the multivariate Gaussian probability distribution function

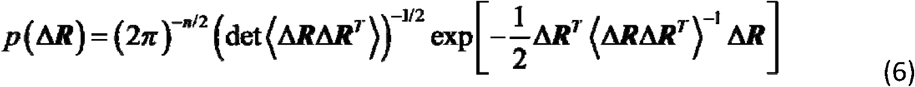

where n is the dimension represented by the number of components of Δ***R*** having n entries which is 3 for a single residue and 6 for a pair. ⟨Δ***R***Δ***R***^***T***^⟩ is the anisotropic covariance matrix, det represents the determinant of the covariance matrix and the superscript t denotes the transpose of the matrix. The angular brackets denote averaging over the full trajectory. In the most general treatment of the protein, Δ***R*** is a 3Nx1 vector, N denoting the number of residues. In this case, *p* (Δ***R***) is a scalar function showing the probability of the state of the protein corresponding to Δ***R***. When a pair of residues is considered only, the effects of the remaining N-2 residues are integrated out from Eq. 6. In this case, ⟨Δ***R***Δ***R***^***T***^⟩ becomes a 6×6 matrix where each of its elements is an average obtained from the molecular dynamics trajectory in the presence of the N-2 residues. Thus, the effects of all the N-2 residues are indirectly present in the 6×6 covriance matrix ⟨Δ***R***Δ***R***^***T***^ ⟩. When Eq. 6 is substituted into Eq. 1 and the average is taken over the full trajectory the following multivariate Gaussian approximation is obtained for the MI between residues i and j^45^

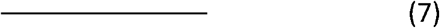

Recent findings demonstrate that convergence of MI values depend strongly on the size of the protein^52^ and the MI predictions of the Gaussian approximation using the multivariate Gaussian model accurately captured mutual information with trajectories of 5 ns for Ubiquitin and 350 ns for the 320 residue long PLpro. Comparison of the MI values from 100 and 300 ns trajectory lengths of the present study showed minimal difference, suggesting convergence of MI for the multivariate Gaussian model for the 191 residue long pyrin.

An alternative approach to mutual information is to use normalized mutual information, NMI defined as: *NMI* (*i, j* = *MI* (*i, j*) / min (*S* (*i*), *S* ( *j*)) where *S* (*i*) and *S* ( *j*) are the entropies of residues i and j. While such a normalization constrains NMI to vary between 0 and 1, it normalizes the raw MI value by the minimum entropy between the variables and may downplay the values of MI especially in allosteric contributions relative to nearby direct contact interactions. Therefore, we report only the raw MI values in the present work.

## Results

### Cavity SASA’s

A closer inspection of the MD simulations of the WT and the six pyrin mutants revealed that the cavity between two loops; one from residue 672 to 680 and the other loop from residue 695 to 697, changed drastically in association with the mutants. In Figure 2. there’s a clear drop in the cavity area of the M694V mutation, which is the most severe form of the disease.

**Figure 2.**
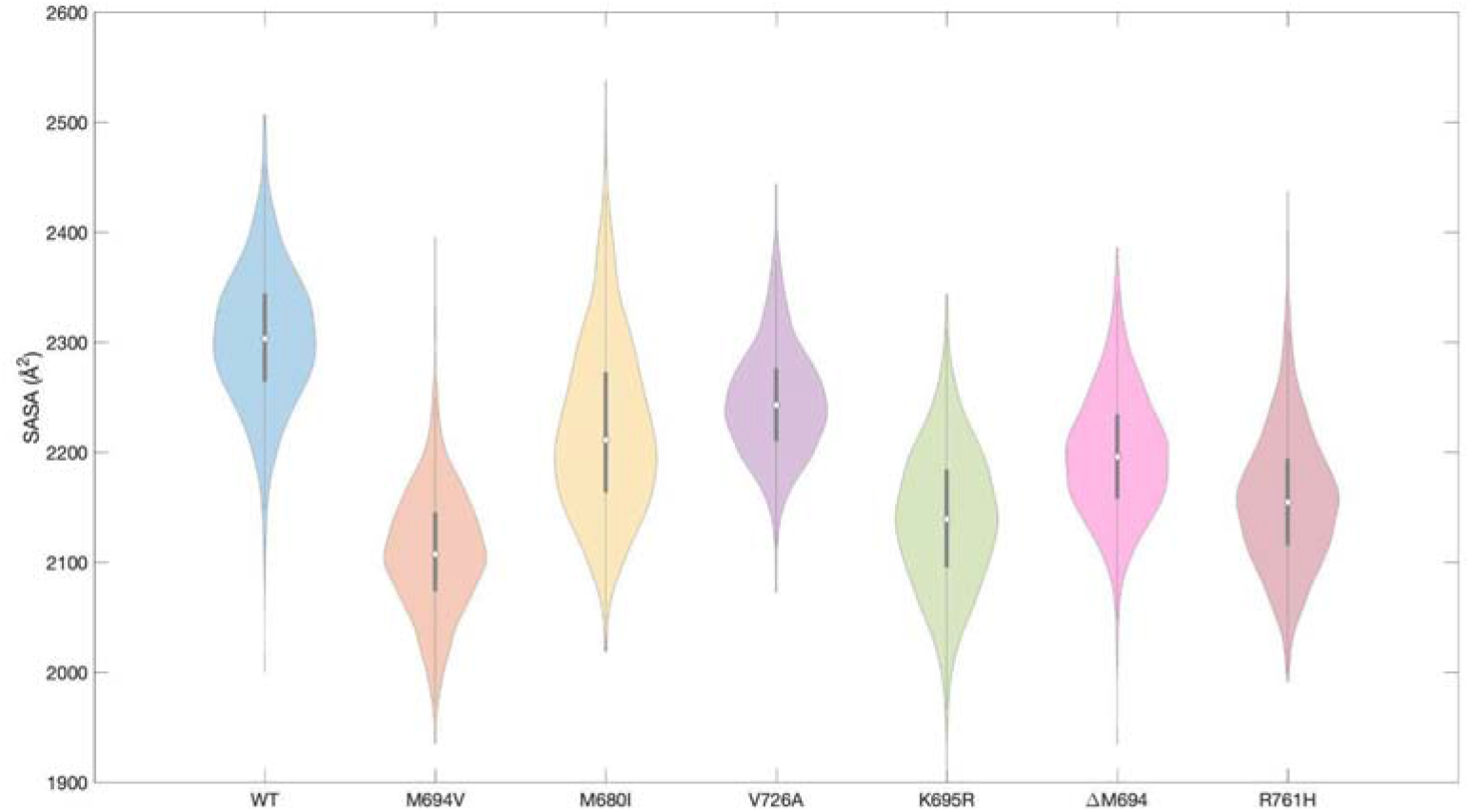
Violin plot of the solvent accessible surface area of the cavity between two loops (loop1: residue 671-683, loop2: residue 695-698) in SPRY domain from the MD simulations of WT, M694V, M680I, V726A, K695R, ΔM694, and R761H mutations.

### K_D_ values from SMD simulations

We performed 10 replicates of SMD simulations for the SPRY and actin complex for the WT and mutants of the SPRY domain to estimate how the K_D_ values change upon mutations. We obtained the pulling force at each time step from the SMD output files and calculated the K_D_ values by using Eq. 3. In Figure 3. we plotted the mean force vs mean distance values from replicate simulations. The average K_D_ value for WT is lower than the mutants, meaning the binding of the WT to actin is stronger. The binding strength from the strongest to the weakest can be ordered as WT (K_D_ = 7.54 pM) > M680I (K_D_ = 12.90 pM) > V726A (K_D_ = 15.10 pM) > R761H (K_D_ = 24.95 pM) > K695R (K_D_ = 47.80 pM) > M694V (K_D_ = 0.25 nM) > ΔM694 (K_D_= 227.35 nM). The mean rupture distances considered for the affinity calculation were selected on the plots.

**Figure 3.**
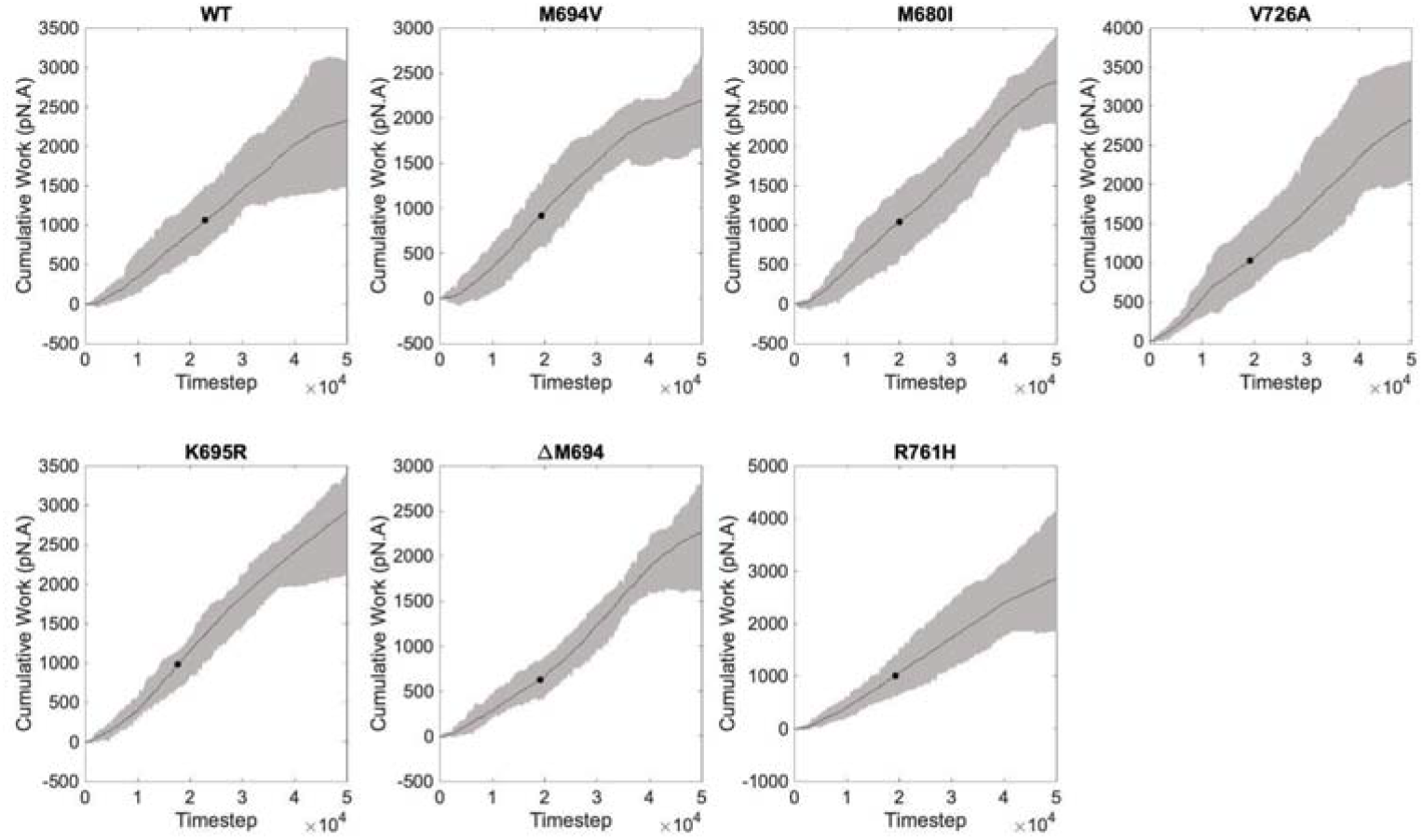
Cumulative work analysis from the replicates of SMD simulations of the WT, M694V, M680I, V726A, K695R, ΔM694, and R761H mutants of Pyrin-SPRY with actin. The shaded region between the maximum and minimum cumulative work curves represents the range of variability observed in the data. The solid line represents the mean cumulative work. The mean rupture point was shown with a dot on the individual plots. The mean rupture distance for each is 11.21 Å, 9.40 A, 9.77 A, 9.41 A, 8.66 Å, 9.38 Å, and 9.40 Å respectively. The rupture distance selection was explained in the methods section.

Our analysis revealed a significant association between low cavity solvent accessible surface areas (SASA) and diminished actin binding. Specifically, mutants with lower cavity SASA exhibited a reduced affinity for actin, suggesting that cavity volume may play a crucial role in mediating the binding interactions between the SPRY domain and actin.

### Mutual information profiles

The findings of our study are summarized in Figure 4, illustrating the mutual information (MI) profile shifts for the four Pyrin mutants. The M694V mutation (upper left panel) exhibits the most substantial negative MI profile shift, followed by the M680I mutation (upper right panel), the V726A mutation (lower left panel), and the K695R mutation with a negligible negative shift (lower right panel). We used the same scale for the abscissae of all panels to facilitate comparison.

**Figure 4.**
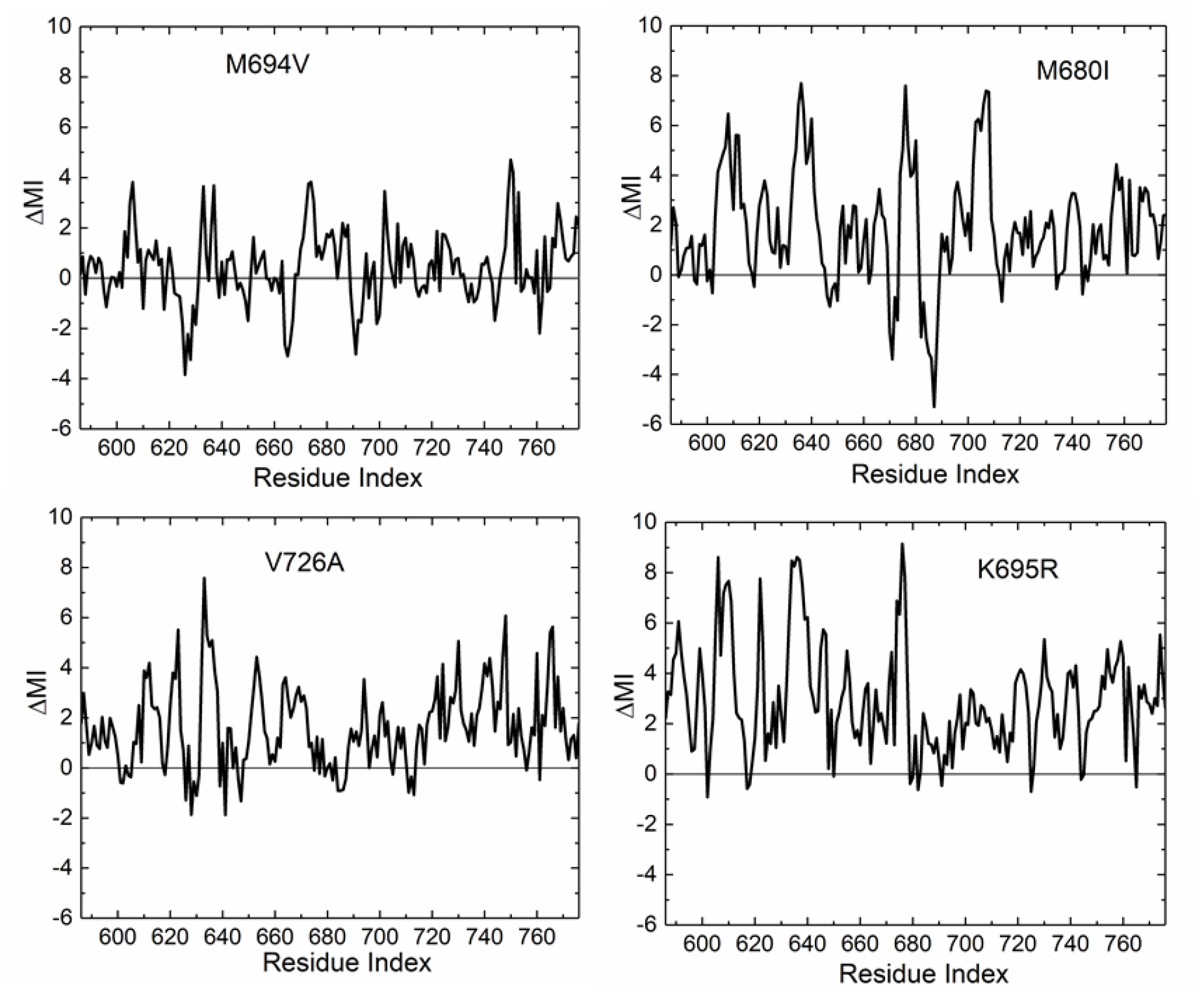
Mutual information profile shifts for the four mutants. Δ*MI* values are calculated using Eq. 2.

An inspection of Figure 4 shows that the negative MI profile shifts are ordered as M694V>M680I>V726A>K695R, and positive shifts are ordered as K695R>M680I>V726A>M694V mutations.

A closer inspection of the panels in Figure 4 provides detailed insights into the presence of both negative and positive MI shifts. When searching for criteria to predict disease severity due to mutations from mutual information (MI) profile changes, focusing on negative changes in MI may be more informative than K_D_ values for the SPRY domain and actin due to the following reasons:

i. For the example of Pyrin, negative changes in MI reflect a decrease in the correlation or interaction strength between residue pairs compared to the WT. These negative changes often indicate the disruption or loss of interactions that have been evolutionarily conserved for maintaining protein structure and function. Such disruptions are more likely to have deleterious effects on protein stability, dynamics, and function, potentially leading to the known disease phenotypes.
ii. On the other hand, increases in MI can result from various factors, including compensatory changes, emergence of new interactions, or alterations in protein dynamics. While increases in MI may sometimes indicate adaptive responses to mutations, they can also be misleading, as they do not necessarily correlate with improvements in protein properties or function. As discussed earlier, increases in MI may not always lead to better protein stability or function and may even have neutral or detrimental effects.
iii. K_D_ values may change for different ligands of the SPRY domain, but the MI shifts are associated primarily with the variations of the SPRY domain of Pyrin protein.

Therefore, focusing on negative changes in MI provides a more direct and biologically relevant criterion for assessing the impact of mutations on protein stability and function. By identifying regions of the protein structure where interactions are lost or disrupted due to mutations, we can prioritize these regions for further investigation as potential determinants of protein function and genotype-phenotype relationship affecting the disease severity.To provide a comprehensive understanding of how mutation affects mutual information across the protein, Figure 5 is presented. This figure illustrates the top 1% of MI gains (depicted as red dots) and the top 1% of MI losses (depicted as black dots) for residue pairs in the four mutants. While the red dots are randomly distributed, showing no consistent patterns of increased contacts in the protein, the black dots reveal distinct patterns of decreased correlations, unique to each protein. Notably, the central region spanning residues 664 to 694 exhibits the most significant mutual information losses across all four mutants. Additionally, smaller clusters of black dots outside the central area demonstrate distinct patterns, indicating loss of mutual information at different parts of the protein depending on the type of mutation.

**Figure 5.**
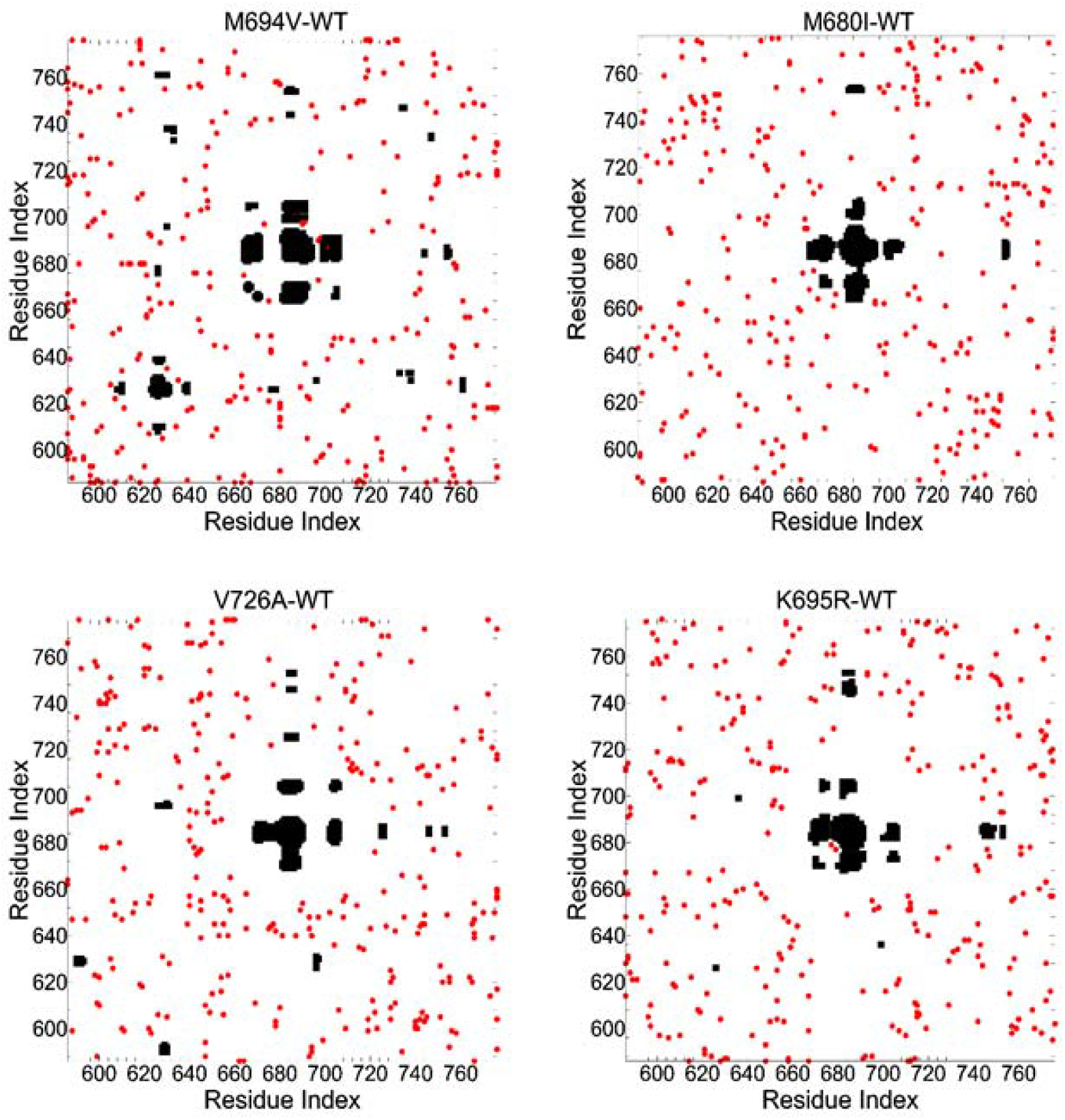
Regions showing Δ*MI* values for the four mutants. Black points show the top 1% of the negative Δ*MI* values. Red points show the top 1% of positive Δ*MI* values.

The central region depicted in Figure 5 features a long beta-hairpin motif, highlighted in blue in Figure 6. The two residues, M694 and A664, are positioned at the ends of the two antiparallel beta strands, while M680 is located at the apex of the long loop. The antiparallel beta strands constitute the base of the binding cavity, to which the D-loop of beta-actin binds. All of the mutations studied here alter the size of the binding cavity, which is the most important factor in determining the function of Pyrin. The three residues shown have significant importance for the function and stability of the protein. The function is controlled by the distance between M680 and M694, which is a measure of the size of the binding cavity. A664, on the other hand, plays a strong role in stability changes with mutation. In the WT, it makes a hydrogen bond with M693 in approximately 94% of total trajectory points whereas this number falls to 1% for the M694V mutation but is not affected much for the other mutations (88% for M680I, 81% for V726A and 83% for K695R).

**Figure 6.**
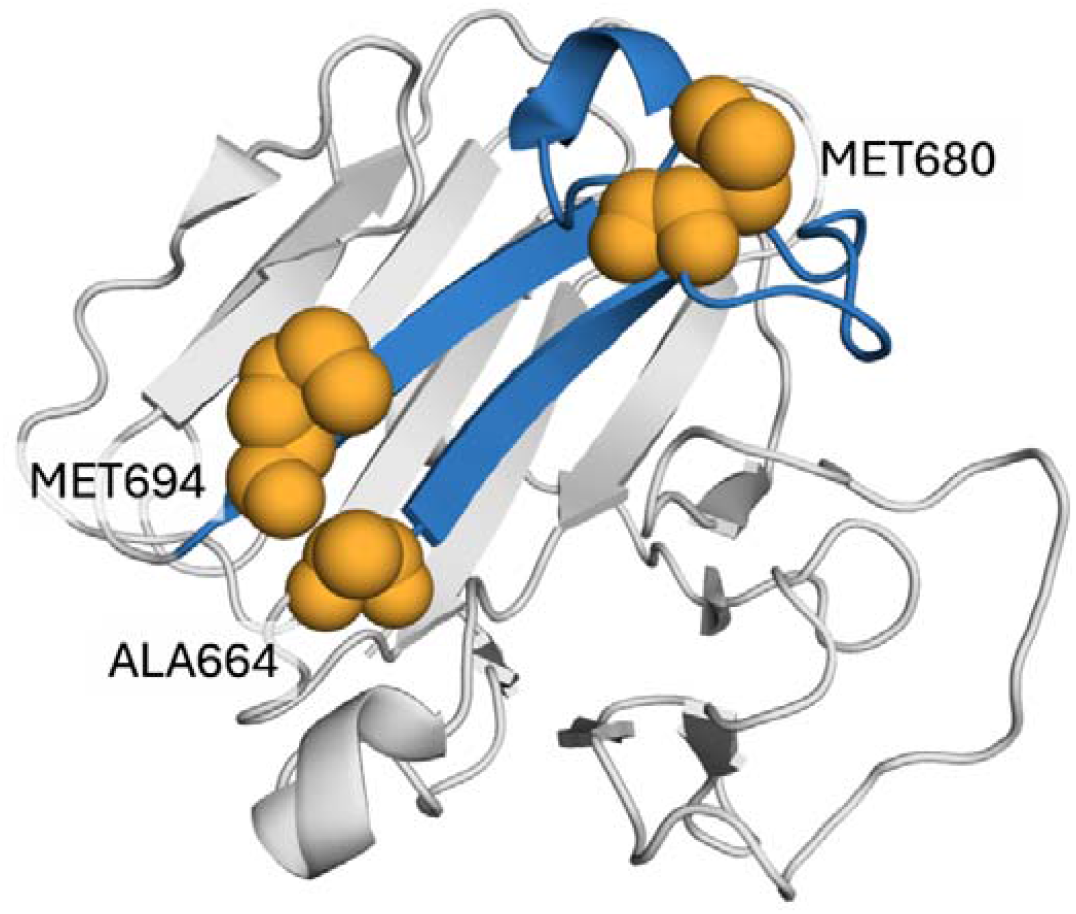
The three-dimensional structure of the SPRY domain of Pyrin. The long beta-hairpin motif is highlighted in blue and corresponds to the central large black dot clusters of Figure 5.

#### 1. Negative MI shifts of M694V

The M694V mutation induces a substantial decrease in pair correlations in one part of the protein followed by a compensatory decrease of pair distances in another part, which affects the function. The central cluster in the upper left panel of Figure 5 is the region of major MI decrease. This cluster is coupled to the smaller clusters, specifically those containing residues K625-R628 and T760-G763 shown in the same panel. Despite being spatially distant, with a distance of 20.9 Å between residues 694 and K625, the mutation at 694 induces a decorrelation effect between residue K625 and several other residues. This raises the question: How does the mutation at residue 694 lead to the decorrelation of residue K625 with the surrounding protein structure? An inspection of the crystal structure shows that there is a path of spatially and covalently contacting residues between K625 and 694 that all lose correlations. The sequence of events leading to this loss of interactions may be described as follows: The mutation leads to a decrease ΔMI=-0.135 between 694 and M693. When M694 is mutated to V694, the hydrophobic interaction between V694 and I692 is lost because the CH3 group of Val rotates in such a way that (i) it starts to interact with K695 and (ii) it approaches E698, and in fact pushes against it. These interactions result in changes in interatomic distances in the neighborhood of residue 694. The probability distribution function, pdf, of the distance between alpha carbons of 694 and E698, *d* (694, 698), is shown with the solid curve in the left panel of Figure 7. The pdf for the WT exhibits three distinct peaks. The left peak corresponds to the case when the hydrogen bond with E698 dominates. The right peak corresponds to the hydrophobic interaction of M694 with I692. This hydrophobic interaction is enhanced by the presence of the two hydrophobic residues M693 and I666.

**Figure 7.**
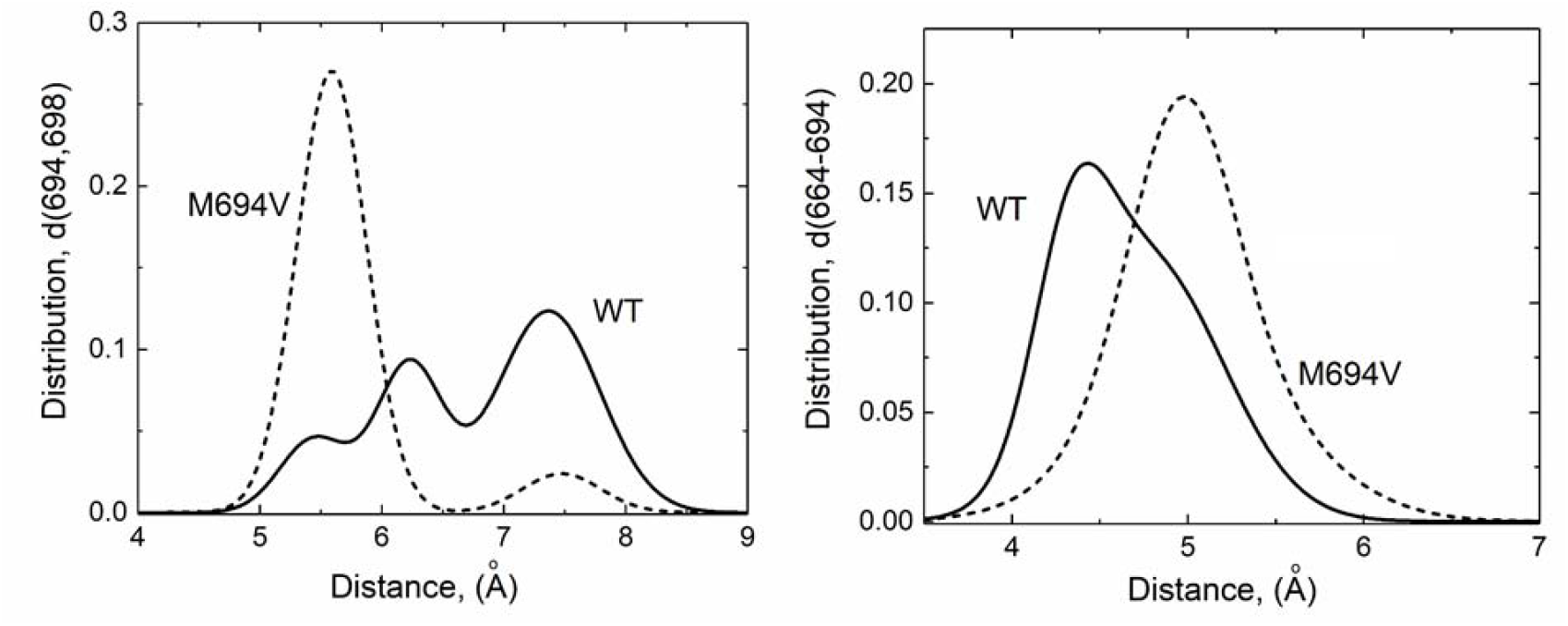
(Left panel) Pdf’s for *d* (694, 698) for WT and the mutant. (Right panel) Pdf’s for *d* (664, 694) for WT and the mutant. The distance between alpha carbons of A664 and 694 is larger in the mutant. In the right panel, the pdf of the WT is skewed towards smaller distances whereas that of M694V is symmetric and at larger distances. Solid line is for the WT, dashed line for the mutant in both panels.

For the mutant, shown with the dashed line in the left panel of Figure 7, there is a dominant peak at short distances corresponding to the interaction of V694 with E698 and K695 and a negligibly small peak at a larger distance. This steric confinement of V694 to the close neighborhood of K695 in the mutant leads to an increase in MI of 0.216 between V694 and K695 and an increase of 0.077 between V694 and E698. When V694 in the mutant moves towards K695 and E698, it moves away from M693 and A664. The increase of the distance between 694 and 664 is shown in the right panel of Figure 7. The pdf’s for the other three mutants are affected to a much lesser extent. This leads to a loss of MI, −0.088, between V694 and A664 and to a loss between V694 and A664. This loss is the major effect of mutation and weakens the interaction between two neighboring beta strands on which V694 and A664 are situated. In general, interactions of 694 with several other residues are lost, although to a lesser extent, as indicated by the negative peaks in Figure 4. The loss of MI progresses from A664 further into the protein as may be seen from the left panel of Figure 8 where the distance between A664 and T760 is increased and the MI between them is decreased. This also affects the neighboring pair R761-K765, shown in the right panel of Figure 8, as a result of which the electrostatic bond between the pair is broken. The loss of the electrostatic interaction and the resulting decrease of MI decreases the stability of the turn on which the pair is located. Because of the destabilizing events mentioned up to here, the corresponding parts of the protein become more flexible. This change in flexibility, specifically in the turns and between neighboring beta strands, makes the mutant vulnerable to interactions that do not exist in the wild type.

**Figure 8.**
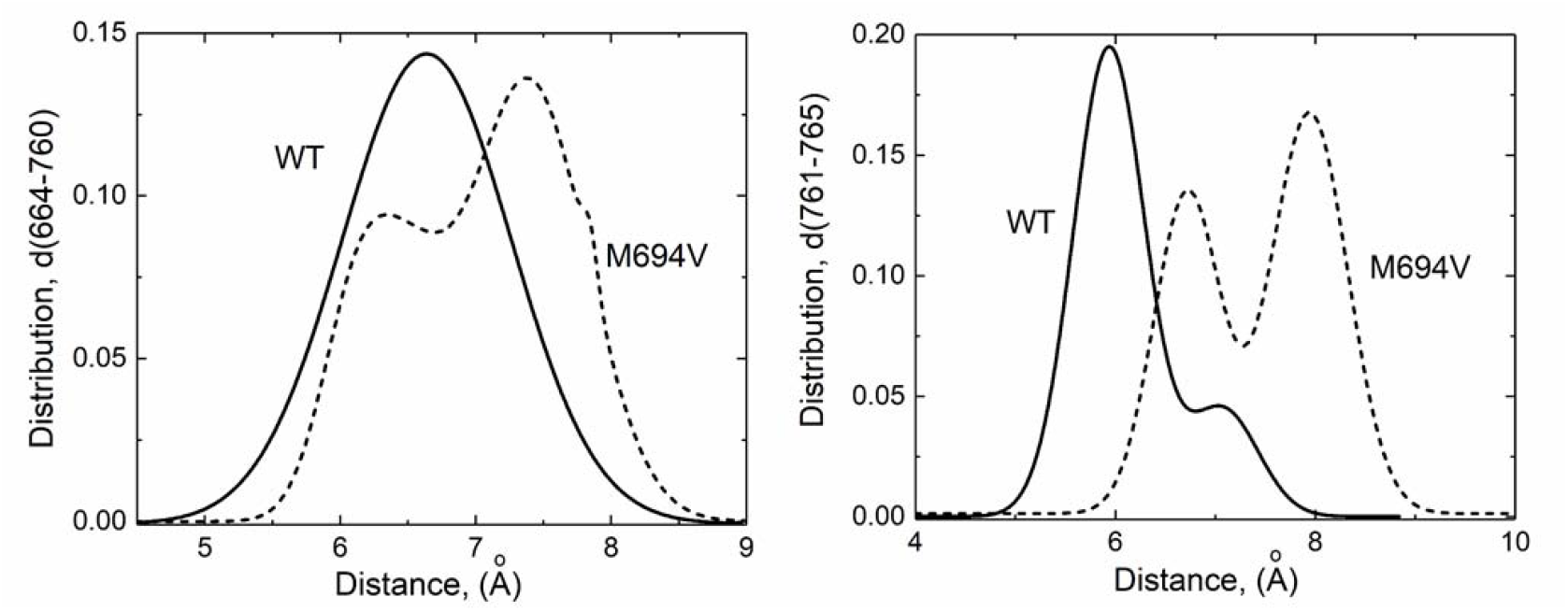
(Left panel) Pdf’s for *d* (664, 760) for WT and mutant. The distance between alpha carbons of A664 and T760 is larger in the mutant. Solid line is for the WT, dashed line for the mutant. (Right panel) Pdf’s for *d* (761, 765) for WT and mutant. The distance between the alpha carbons of R761 and K765 is larger in the mutant. The solid line is for the WT, dashed line is for the mutant for both panels.

The interactions of R761 are crucial in the behavior of the WT and the mutant. MI is lost in the loop spanning the residues P758 and T767. Similarly, MI is lost between R761 and the large loop spanning residues N624 to F636 which contains the residue E627 but the interactions between R761 and S638, K662, M680, S703, K712, G750, P754, P758, and N766 are increased.

Although E627 and R761 make an electrostatic bond through their exterior Oxygen and Nitrogen atoms in the WT structure, their alpha carbons are intercalated by an intermediate segment. In the mutant, R761 becomes highly mobile and flexible, due to the instability of the loop bordered by R761 and K765 as indicated by the pdf’s in the right panel of Figure 8. This enhanced mobility removes various interactions and introduces newer ones. Firstly, the interactions of R761 with E627 are affected. The distribution of the distance between the alpha carbons of E627 and R761, Figure 9, left panel, shows that these two residues are closer to each other in the M694V mutant compared to the WT. The distribution of the alpha carbon distances is Gaussian and symmetric in the WT structure, but it becomes skewed in the mutant, preferring shorter distances, resulting in steric clashes between the atoms of the two residues. The right panel shows the distribution of the electrostatic bond length between O and N of E627 and R761, respectively. The sharp peak in the WT curve indicates the stable electrostatic bond in the WT, whereas this is lost in the mutant and the electrostatic bond length is compressed.

**Figure 9.**
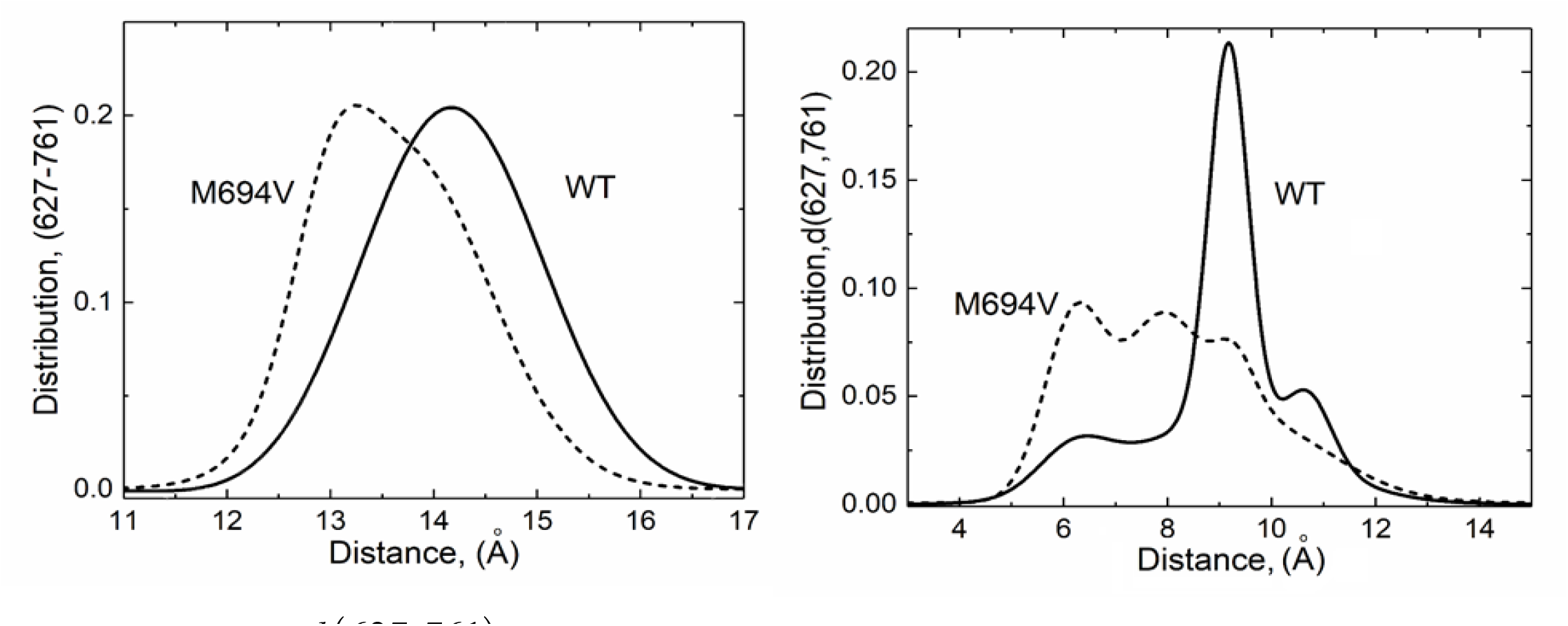
Pdf’s for *d* (627, 761) for WT and mutant. The left panel is for the distance between alpha carbons, and the right panel for the length of the electrostatic bond. In both cases, the distance between 627 and 761 is smaller in the mutant. The solid line is for the WT, the dashed line is for the mutant.

The change in the interaction of E627 and R761 is important. The E627-R761 pair acts as a hinge that introduces flexibility to the system, leading to the decrease in the distances between the following pairs: K662-N766, R761-L622, M680-R761, S638-M680. These are shown in Figure 10. Among these, the decrease in the distance S638-M680 results from the increased flexibility of the long loop spanning residues G623 to P633 which arises from the loss of MI between V694 and A664. Thus, the path V694-A664-G623-M680 controls the approach of M680 to M694. A more detailed account of the path and MI changes is shown in Table 2.

**Table 2:** The decorrelation path in M694V.

| Residue Pair | M694-M693 | M693-A664 | A664-K765 | K765-R761 | R761-E627 | E627-K625 |
| --- | --- | --- | --- | --- | --- | --- |
| $\Delta MI$ | -0.135 | -0.117 | -0.010 | -0.194 | -0.036 | -0.089 |

**Figure 10.**
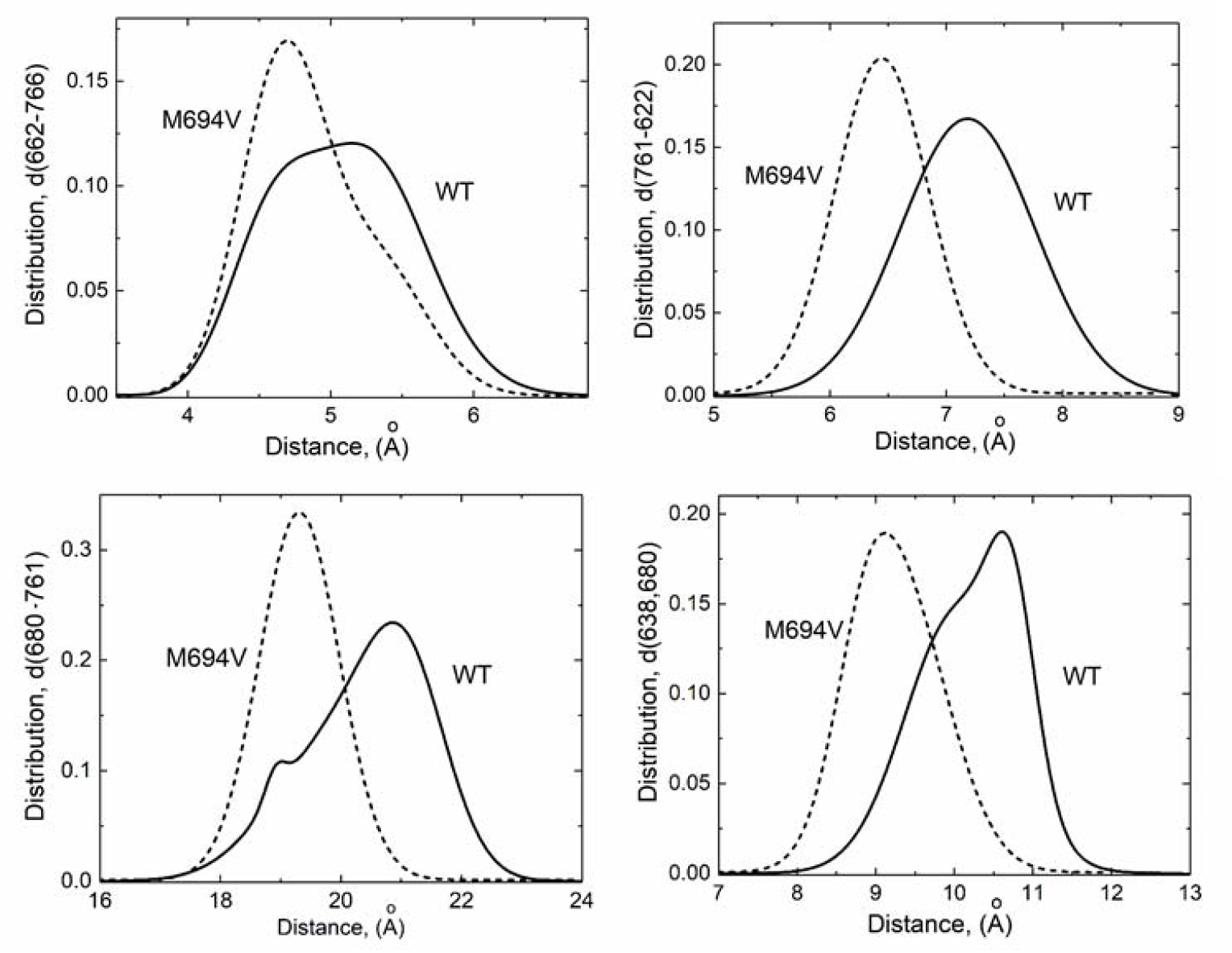
The decrease in the distance between various pairs in the mutant. The solid line is for the WT, the dashed line is for the mutant.

**Figure 11.**
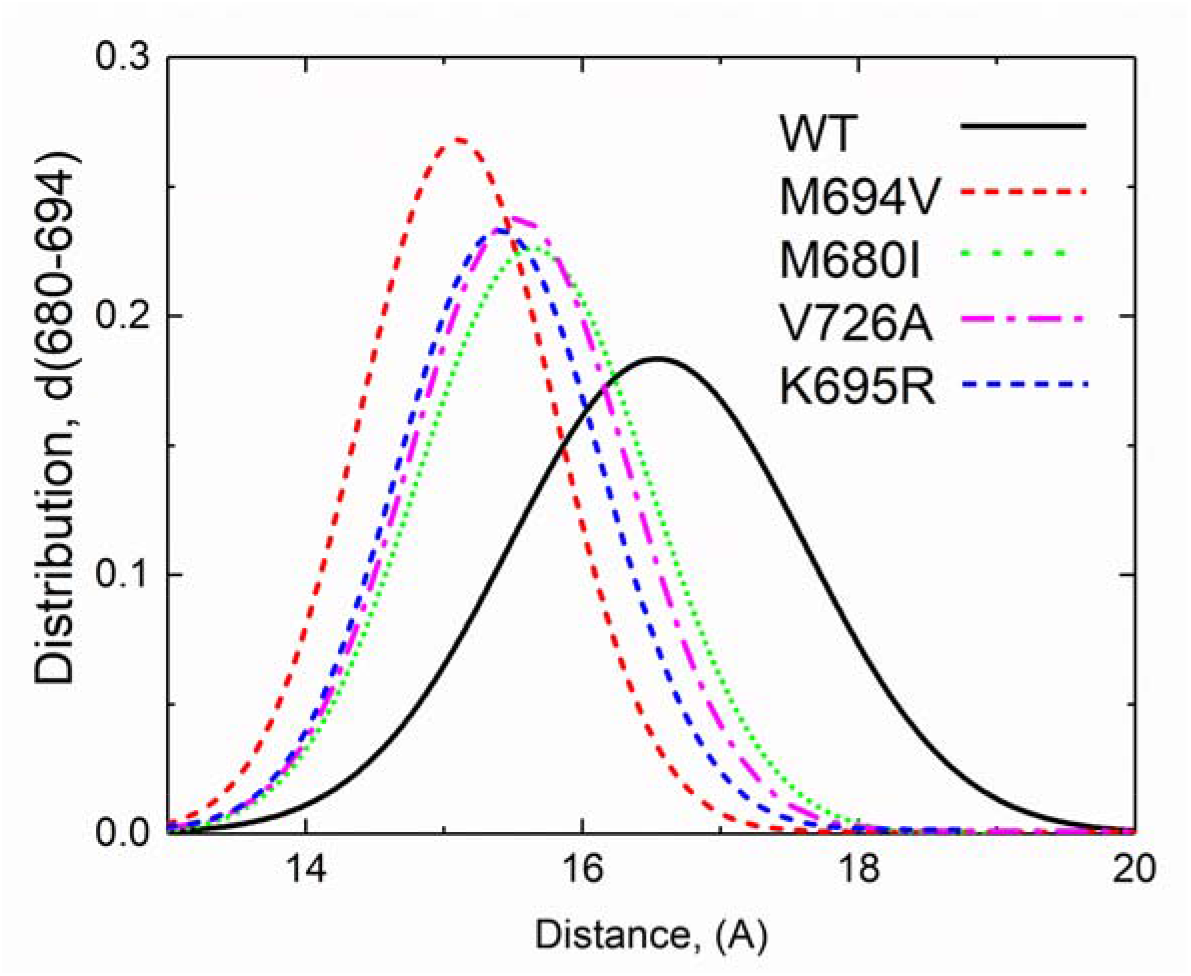
Change in distance between 680 and 694 in WT (black solid line) and the mutants (dashed lines). The curve for the R761H is in between V726A and M680I and is not shown.

Roughly assuming pairwise additivity, the loss of interaction along the path shown in Table 2 accounts for approximately 15% of the cumulative loss of K625 interactions. As discussed further in the Discussion section, it is important to note that contacting residues in a native structure is typically at a minimum energy state and maximally correlated due to evolutionary optimization. Any perturbation to the structure, such as the mutation discussed above, is expected to lead to a decrease in correlations, as demonstrated in this example. The decrease in the distance between M680 and M694, which affects the binding of Actin to the binding pocket of Pyrin is a consequence of these changes.

#### 2. M680I mutation causes a decrease in the distance between M680 and the neighboring F636-C639 loop

Changes in the conformation of the long hairpin beta motif resulting from M680I mutation result from two compensatory motions. The first is the increase in the 680 and N686 distance. This increase results from the substitution of Ile for Met at 680 which leads to the loss of several hydrogen bonds that stabilize the turn on which 680 and N686 lie. MI between the pair M680-N686 decreases. For the other mutants, V726A and K695R, this decrease is insignificant and the change is positive for the M694V mutant. In more detail, S683 and N686 on the 680-N686 loop form two hydrogen bonds in the WT crystal structure which are observed in 71% of the time points in the molecular dynamics trajectories. In the M680I mutant, this frequency falls to 55%. In the other three mutants, M694V, V726A, and K695R they are observed to be 70.0%, 73.8%, and 69%, respectively. Decorrelation of the 680-loop resulting from hydrogen bond loss appears to be the significant factor that affects the conformation of the long beta-hairpin loop motive in the M680I mutant while the motive in the other mutants is not affected. To compensate for the increase in the 680 and N686 distances, the neighboring loop distances, i.e., the 680-F636 and 680-C639 distances are decreased (See Figure S-1). In the crystal structure of Pyrin, there is a Pi-Sulfur bond between M680 and F636 and a hydrophobic interaction between M680 and C639. Decorrelation of the 680-N686 interaction results in a significant decrease in the length of the 680-F636 and 680-C639 bonds. These changes distort the geometry of the long hairpin beta motif on which the D-loop of beta-actin binds

#### 3. Allosteric Interactions in Pyrin. Propagation of effects in V726A

The relatively less penetrant variant V726A is not located in the protein binding cavity of the SPRY domain. The distance between the alpha carbons of V726 and M680 in the crystal structure is 26.2Å. Replacing Valine of residue 726 with Alanine removes two CH3 groups and inserts one CH3 which results in a decrease in steric interactions around 726, notably in correlations of A726 with G727. This decorrelation results from enhanced covalent angle changes between A726 and G727 which make the junction of the two act like a hinge. This leads to a decrease of MI along a path between A726 and M680 (See Figure S-2) and an increase of the distance between A726 and S675 (See left panel of Figure S-3) and to their decorrelation which triggers the decorrelation and increase of distance between S675 and 680. (See right panel of Figure S-3).

#### 4. K695R and similarity to M694V

The mechanism of this mutation is similar to that of the M694V mutation with milder consequences. In the WT structure, K695 makes three hydrogen bonds with E698 but in the mutated structure, R695 makes five hydrogen bonds with E698. The result is a decrease in the distance between the two in the mutant (See right panel of Figure S-4). This change introduces decorrelations into the system similar to those in M694V but weaker. The decreases in MI values between residue 694 and the rest of M694V follow a similar trend with M694V, but the decreases being weaker in the former (see Figure S-5).

#### 5. The case of R761H

While the role of R761 is consequential in the case of the M694V mutant, as discussed above, its role in the R761H mutant is not as effective. Like V726, R761 is outside the binding cavity. In the crystal structure, R761 makes an electrostatic bond with E627, a hydrogen bond with L622, and hydrogen bonds with G764, K764, and N766. Hydrogen bonds with the latter three stabilize the loop that contains R761. Mutation into His keeps the three stabilizing loop hydrogen bonds but loses interaction with E627, which is not directly consequential in the interactions of the binding cavity residues. The effect of the mutation changes the mobility of the loop on which 761 lies and of the long loop on which E627 lies. While R761 lies on the decorrelation path discussed above concerning M694V, it is less penetrant because M694 lies directly on the long beta-hairpin motif but R761 does not. Therefore, whatever role R761 plays in the distortion of the binding area is indirect and allosteric. The MI losses due to the R761H mutation are shown in Figure S-6 where the dominant loss is an allosteric loss of interaction for the distant pair, 761-694.

#### 6. Change in the distance between 680 and 694 for all mutants

The functionally important result of the studied mutations is the decrease in the size of the binding cavity, as evidenced by the reduced distance between residues 680 and 694, serving as a linear indicator of the area.

#### 7. MI losses resulting from M694 deletion

The focus of the present study is MI losses upon missense mutations. Nevertheless, we also evaluated the MI losses for the deletion of M694 (ΔM694) as being the most severe form of the variants in the SPRY domain resulting in an autosomal dominantly inherited phenotype in comparison to all other recessively inherited mutants. Removal of a residue led to extreme loss of MI as can be seen in Figure S-7 and to a decrease in the binding area and a change in the distance distribution as can be seen in Figures 2 and S-8 respectively.

## Discussion

In this study, we report a novel method to assess the pathogenicity of Pyrin mutations leading to the FMF phenotype and/or its severity by using the multivariate Gaussian representation of MI as a measure to quantify the correlation between residue motions or fluctuations. This method involves forming six-dimensional vectors for each pair of residues i and j (i.e., Δ*x*_*i*_, Δ*y*_*i*_, Δ*z*_*i*_, Δ*x*_*j*_, Δ*y*_*j*_, Δ*z*_*j*_) and calculating the covariance matrix from the molecular dynamics trajectory data. The Pearson correlation coefficient, PCC, is another method for measuring dependencies between variables, and both the multivariate Gaussian and the Pearson correlation are inherently linear. However, our approach using the six-dimensional multivariate Gaussian model is more comprehensive because it considers the full covariance structure, including cross-terms such as ⟨Δ*x*_*i*_ Δ*y*_*j*_⟩ This enables us to capture the joint dependencies across multiple dimensions, thus providing a richer and more detailed understanding of the correlations between residue fluctuations. This is crucial for protein structure analysis, where anisotropic interactions need to be included.

Our analysis using MI revealed distinct patterns for the FMF-associated mutations in the pyrin protein. Negative MI shifts in Figure 4, signifying a loss of correlation between residue fluctuations, were observed along well-defined pathways in the protein structure as clearly seen in Figure 5. These pathways likely represent functionally important communication networks disrupted by the mutations, potentially leading to protein dysfunction and disease development. In contrast, positive MI shifts, indicating an increase in correlation between residues, appeared as random points scattered across the structure, see Figure 5. While positive shifts can, in some instances, introduce detrimental rigidities or hinder necessary conformational changes, their seemingly random distribution suggests a lack of well-defined patterns in this context. These might not directly influence the protein’s core functional motions. They could represent incidental increases in correlation arising from local structural changes caused by the mutation, which don’t necessarily contribute to the disease pathology. The observation that negative shifts exhibit clearly defined residue contact paths highlights their potential role in disrupting critical communication pathways within the pyrin protein.

We have used FMF-associated Pyrin mutations to test the performance of MI measurements in the prediction of pathogenicity since most of the available computational tools used to predict the pathogenicity of the known missense mutations were not helpful for the case of the most pathogenic Pyrin variants. Our study reveals that the negative MI profile shifts of the mutants are ordered as M694V>M680I>V726A>K695R in the same order as the pathogenicity and disease severity in FMF patients.

The Cavity SASA calculation quantifies the volume of the ligand-binding cavity within the SPRY domain of Pyrin protein. Our finding that the mutation decreases this volume provides structural context for the functional changes observed in association with the actin ligand. A smaller cavity might hinder the proper binding of the actin D-loop. The steered MD simulations and resulting K_D_ values directly assess the functional consequence of the mutation on actin binding. The decrease in K_D_ indicates a weaker interaction between the mutated pyrin and actin, supporting the notion that the mutation disrupts the binding process. The MI analysis reveals a decrease in the correlated fluctuations of residues bordering the cavity upon mutation. This suggests that the mutation disrupts the communication pathways within the protein, potentially affecting the dynamics of the cavity and its ability to bind actin. By integrating the structural changes (Cavity SASA) with the functional consequences (K_D_) and the underlying communication network disruptions (MI), we gain a deeper understanding of how the mutation impacts the pyrin-actin interaction.

Although the importance of the SPRY domain and the positions of Met694 and Met680 amino acids at the entrance of the shallow cavity have long been known in the pathogenesis of FMF, no previous study investigated the correlation between the variant-related changes in the cavity area and the disease severity. If we ignore the very rare and dominantly inherited variant of ΔM694, the M694V mutation has the highest Pras severity score among other missense variants in parallel with clinical severity findings^53-56^. Recent studies analyzing the functional consequences of the FMF-associated SPRY domain variants using an in vitro test platform provided evidence supporting the clinical severity findings. The M694V mutation was considered to be the most pathogenic FMF allele causing severe disease. The in vitro diagnostic test distinguished samples collected from the patients with the known FMF mutations (M694V, M694I, M680I, R761H) from the samples from patients with other MEFV mutations considered of uncertain clinical significance (K695R). Within this context, our current work provided very consistent results with clinical experience. Finally, it should be noted that while all-atom molecular dynamics (MD) simulations offer detailed insights into mutation-disease relationships through the characterization of MI profile shifts, their computational cost restricts the analysis to a limited set of proteins.

## Acknowledgment

This project was partially supported by TUBITAK 1004 INFLAM-IST Project (20AG007).

