## Supplemental Figures for "Role of Mutual Information Profile Shifts in Assessing the Pathogenicity of Mutations on Protein Functions: The Case of Pyrin Variants Associated with Familial Mediterranean Fever"

The behavior of M680I, V726A, K695R, and R761H, are referred to in the main text but more detailed information is presented in this Supplementary Material section. The section numbers and section headings here correspond to those of the main text.

**2. M680I mutation causes a decrease in the distance between M680 and the neighboring F636-C639 loop**: Changes in the conformation of the long hairpin beta motif resulting from M680I mutation result from two compensatory motions. The first is the increase in the 680 and N686 distance. This increase results from the substitution of Ile for Met at 680 which results in the loss of several hydrogen bonds that stabilize the turn on which 680 and 686 lie. MI between the pair M680-N686 decreases. For the other mutants, V726A and K695R, this decrease is insignificant and the change is positive for the M694V mutant.

In more detail, S683 and N686 on the 680-686 loop form two hydrogen bonds in the WT crystal structure which are observed in 71% of the time points in the molecular dynamics trajectories. In the M680I mutant this frequency falls to 55%. In the other three mutants, M694V, V726A and K695R they are observed to be 70.0%, 73.8% and 69%, respectively. Decorrelation of the 680-loop resulting from hydrogen bond loss appears to be the significant factor that affects the conformation of the long beta hairpin loop in the mutant M680I mutant while the other mutants are not affected. To compensate for the increase in the 680 and N686 distance, the M680-C636 and M680-C639 distances are decreased, shown in Figure 10. In the crystal structure of Pyrin, there is a Pi-Sulfur bond between M680 and F636 and a hydrophobic interaction between M680 and C639. Decorrelation of the 680-686 interaction results in a concomitant decrease in the length of the 680-636 and 680-639 bonds significantly as shown in Figure S-1. These changes distort the geometry of the long hairpin beta motif where the D-loop of Actin binds.

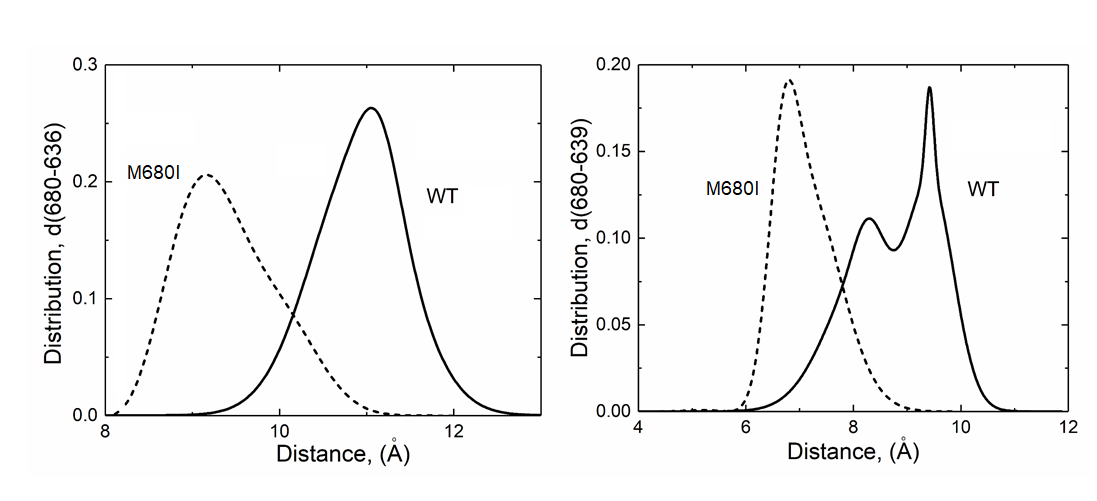

Figure S-1. (Left panel) Distance changes between 680 and 636. (Right panel) Distance changes between 680 and 639. Solid curves are for the WT, dashed curves for mutants for both panels. Other mutants, M694V, V726A and K695R do not show this decrease.

**3. Allosteric Interactions in Pyrin. Propagation of effects in V726A**: The relatively less penetrant variant V726A is not located in the protein binding cavity. Replacing Valine of residue 726 with Alanine removes two CH3 groups and inserts one CH3 which results in a decrease in steric interactions around 726, notably in correlations of A726 with G727. Figure S-11 shows the decrease of MI values between A726 and the residues along the path to M680. The small red circles show the residues along a path between A726 and M680. The mutation of residue V726 to Alanine causes it to strongly decorrelate with the upstream neighbor G727. The decorrelation results from enhanced angle changes between A726 and G727 which makes the junction of the two act like a hinge. This decorrelation effect propagates along the path mentioned but the distances between successive pairs on the path are not affected. However, the decorrelation shown in Figure S-1 with the black circle, A726-S675 results in an increase in distance, shown in the left panel of Figure S-2.

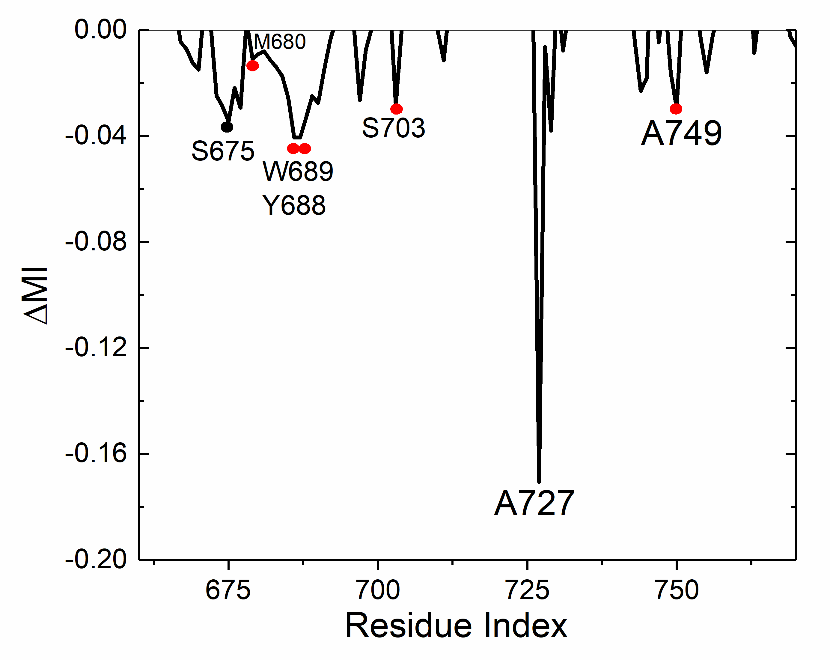

Figure S-2. Residues showing MI decrease with A726 shown with circles. The black circle shows a residue that is not on the path.

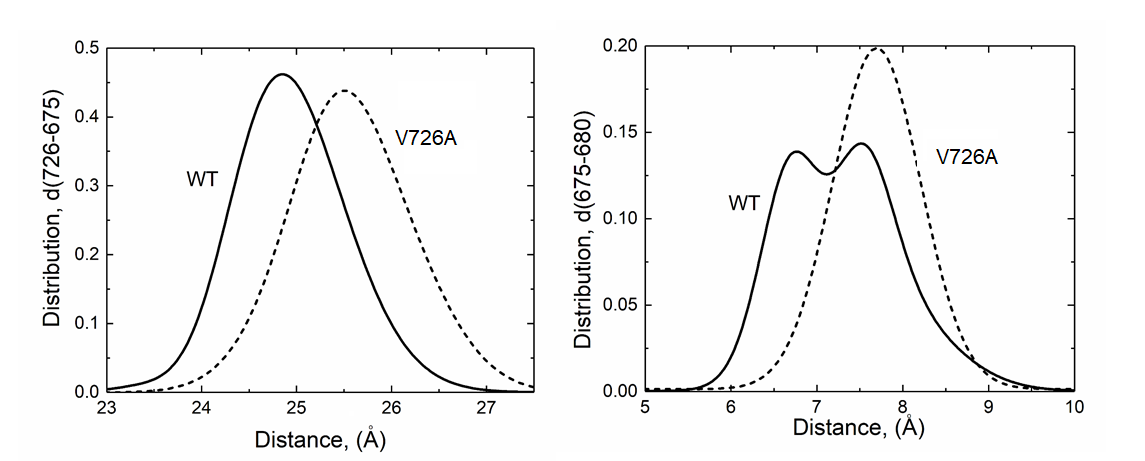
Figure S-3. Change of distance between 726 and 675 (Left panel) and between 675 and 680 (right panel) . Solid line is for the WT, dashed line for the mutant.

The decorrelation between 726 and 675 triggers the decorrelation between 675 and 680 which increases the mean distance between 675 and 680 shown on the right panel of Figure S-2. The distance between 726 and 680 increases as shown in the left panel of Figure S-3.

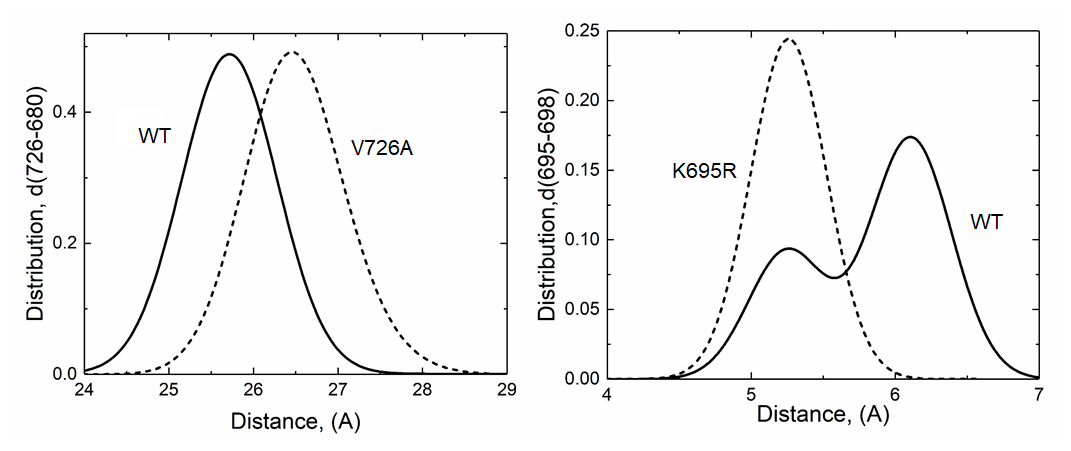

Figure S-4. (Left panel) Distance between V726 and M680 in V726A. (Right panel) Distance between K695 and E698 in K695V. Solid line is for the WT, dashed line for the mutant in both panels.

**4. K695R and similarity to M694V**. The mechanism of this mutation is similar to that of the M694V mutation with milder consequences. In the WT structure, K695 makes three hydrogen bonds with E698 but in the mutated structure R695 makes five hydrogen bonds with E698. The result is a decrease in the distance between the two in the mutant as shown in the right panel of Figure S-4. This change introduces decorrelations into the system similar to those in M694V but weaker. The decreases in MI values between residue 694 and the rest of the protein are shown in Figure S-5.

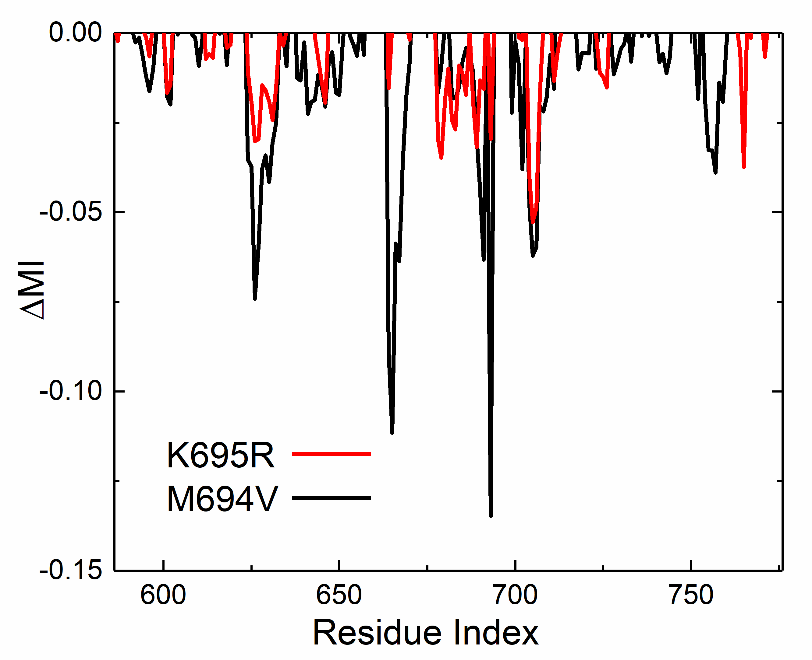

Figure S-5. Comparison of MI decreases for K695R and M694V.

**5. The case of R761H**. R761 is outside the binding cavity and plays an indirect allosteric role in distorting the binding area. The MI losses due to the R761H mutation are shown in Figure S-15 where the dominant loss is an allosteric loss of interaction for the distant pair, 761-694.

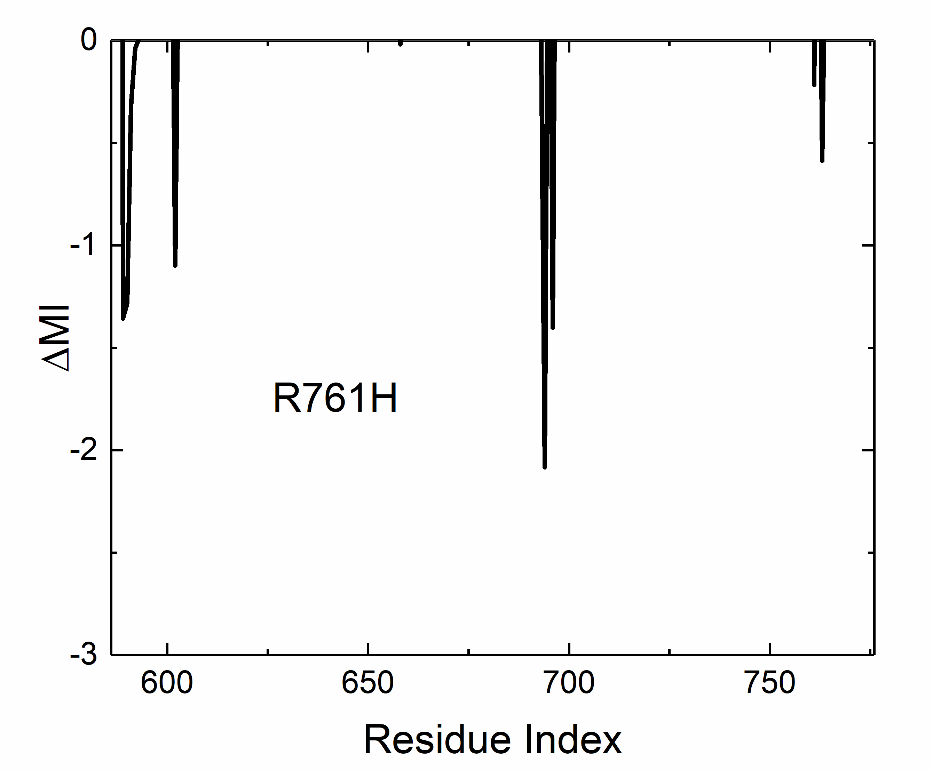

Figure S-6. MI losses due to the R761H mutation.

**7. MI losses resulting from M694 deletion**. MI losses for the deletion of M694 are the most severe form of the variants in the SPRY domain resulting in an autosomal recessively inherited phenotype. Removal of a residue led to extreme loss of MI as can be seen in Figure S-7 and to a decrease in the binding area as can be seen from the decrease of the distance between 680 and 694 in Figure S-8.

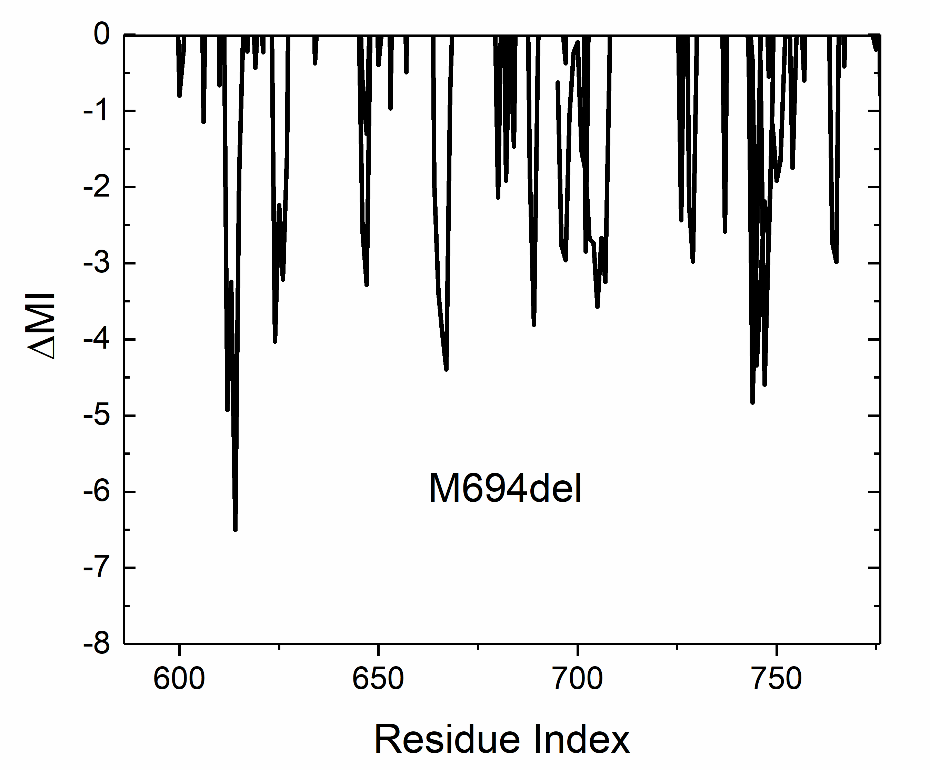

Figure S-7. MI losses in M694del relative to the WT.

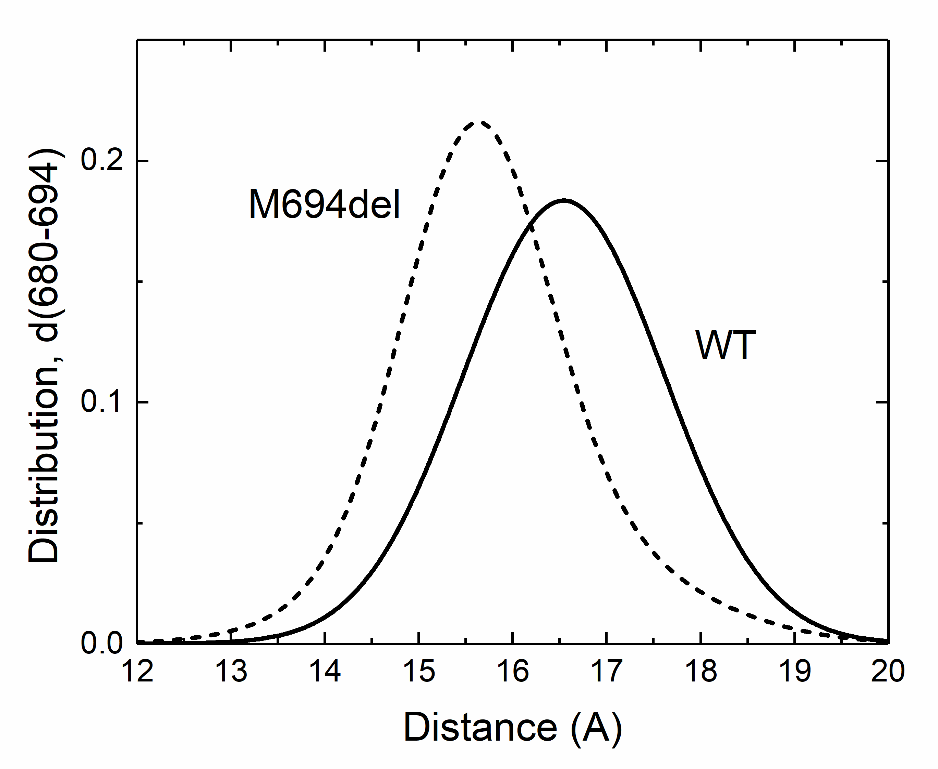

Figure S-8. Change in the distance between 680 and 694. Removal of M694 from the WT brings K695 in its place in the chain. After deleting residue M694 in Pyrin, residue K695 of the original chain becomes the chemically bonded neighbor of M693. Subsequently, all residues beyond K695 are renumbered, with K695 now becoming K694 in the renumbered chain.
